# Development of Potent, Selective cPLA2 Inhibitors for Targeting Neuroinflammation in Alzheimer’s Disease and Other Neurodegenerative Disorders

**DOI:** 10.1101/2025.03.09.641748

**Authors:** Anastasiia V. Sadybekov, Marlon Vincent Duro, Shaowei Wang, Brandon Ebright, Dante Dikeman, Cristelle Hugo, Bilal Ersen Kerman, Qiu-Lan Ma, Antonina L. Nazarova, Arman A. Sadybekov, Isaac Asante, Stan G Louie, Vsevolod Katritch, Hussein N. Yassine

**Affiliations:** Department of Quantitative and Computational Biology, University of Southern California, Los Angeles, CA, USA; Center for New Technologies in Drug Discovery and Development, Bridge Institute, Michelson; Department of Radiology, Keck School of Medicine; Department of Neurology, Keck School of Medicine; Department of Clinical Pharmacy, Mann School of Pharmacy; Department of Ophthalmology, Keck School of Medicine; Michelson Center for Convergent Biosciences, University of Southern California, Los Angeles, CA, USA; Department of Pharmacology and Pharmaceutical Sciences, Mann School of Pharmacy; Center for Personalized Brain Health, University of Southern California, Los Angeles

**Keywords:** cPLA_2_, neuroinflammation, neurodegeneration, Alzheimer’s disease

## Abstract

Chronic neuroinflammation plays a key role in the progression of Alzheimer’s disease (AD), and the cytosolic calcium-dependent phospholipase A2 (cPLA_2_) enzyme is a critical mediator of inflammatory lipid signaling pathways. Here we investigate the therapeutic potential of novel cPLA_2_ inhibitors in modulating neuroinflammation in AD. By leveraging the giga-scale V-SYNTHES 2.0 virtual screening in on-demand chemical space and conducting two rounds of optimization for potency and selectivity, we have identified BRI-50460, achieving an IC_50_ of 0.88 nM in cellular assays of cPLA_2_ activity. *In vivo* studies revealed favorable brain-to-plasma ratios, highlighting the ability of BRI-50460 to penetrate the central nervous system, potentially modulating neuroinflammatory pathways and restoring lipid homeostasis. In cultured astrocytes and neurons derived from human induced pluripotent stem cells, BRI-50460 mitigates the effects of amyloid beta 42 oligomers on cPLA_2_ activation, tau hyperphosphorylation, and synaptic and dendritic reduction. Our results suggest that small molecule inhibitors of the cPLA_2_ enzyme can modulate the downstream inflammatory lipid signaling pathways, offering a promising therapeutic strategy for AD and other neurodegenerative diseases.

## Introduction

The lack of effective treatments for neurodegenerative diseases such as Alzheimer’s disease (AD) underscores the challenge in identifying the critical pathways that influence neurodegeneration. Neuroinflammation is a hallmark of Alzheimer’s disease (AD) and other neurodegenerative disorders. Inflammation-related targets constitute one of the largest growing categories (approximately 16%) in the AD drug development pipeline^1^. Understanding the complex role of neuroinflammation in disease pathogenesis unveils more effective therapeutic approaches and druggable targets. Among the inflammatory pathways driving chronic, unresolved inflammation are pro-inflammatory mediators generated from arachidonic acid (AA) metabolism^2–4^. The enzyme cytosolic phospholipase A_2_ (cPLA_2_) plays a critical role in this process by hydrolyzing membrane phospholipids to release AA, which is subsequently converted to both pro-inflammatory and anti-inflammatory eicosanoids, with greater representation of proinflammatory pathways.

One of the key lipoproteins regulating brain lipid metabolism is apolipoprotein E (ApoE), which exists in three common variants: ApoE2, ApoE3, and ApoE4. The *APOE4* allele is not only the strongest genetic risk factor for late onset AD but also has been shown to exacerbate neuroinflammation^5^. Our group has reported increased cPLA_2_ activity in *APOE4* cellular and animal models, as well as in postmortem brain tissues, contributing to lipid dysregulation and neuronal damage. In this context, cPLA_2_ inhibitors may offer a novel therapeutic approach by addressing both unresolved inflammation and lipid homeostasis. cPLA_2_ catalyzes the hydrolysis of phospholipids, resulting in the production of free fatty acids (FA) and lysophospholipids^6^. As *APOE4* enhances the activation of cPLA_2_ ^17^ in the brain, the ω-6 to ω-3 balance is disrupted. Reducing cPLA_2_ activity realigns the ω-6 to ω-3 balance^8^, suggesting an attractive therapeutic strategy.

The development of selective cPLA_2_ inhibitors has proven challenging due to off-target effects and difficulties with isoform specificity, with the additional challenge of blood-brain barrier (BBB) penetrance. After a decade of developing these compounds, only one has gone through human trials (AK106-001616). A Phase 2 clinical trial explored AK106-001616 in patients with rheumatoid arthritis (RA)^9^. While clinical safety for AK106-001616 has been established in RA patients, its ability to penetrate the BBB is likely to be limited. Therefore, new, specific, and brain-penetrant cPLA2 inhibitors are needed.

In this study, we employed V-SYNTHES 2.0, a novel approach to virtually screen the Enamine REAL (REadily AvailabLe) Space of 36 billion on-demand synthesizable compounds for potential cPLA_2_ inhibitors. Optimization of the initial hits in REAL Space resulted in several lead compounds that merit further development. The most promising candidate BRI-50460 potently and selectively inhibits cPLA_2_ in vitro and in our cellular experiments; it also mitigates the effects of amyloid beta 42 oligomers (Aβ42O) on cPLA_2_ activation, tau hyperphosphorylation, and synaptic and dendritic reduction. These results offer a therapeutic strategy that not only targets the enzyme itself but also modulates the downstream lipid signaling pathways associated with AD pathology.

## Methods

### Drug Discovery using V-SYNTHES 2.0 Platform

#### cPLA_2_ Model Preparation

The prospective screening was performed using a structural model of human cPLA_2_ refined using a ligand-guided approach^10^. A 2.5 Å resolution cPLA_2_ crystal structure that includes a *N*-terminal calcium-dependent lipid-binding/C2 domain, and a catalytic unit was downloaded from PDB (PDBID:1CJY). The crystal structure of cPLA_2_ was converted from PDB coordinates into an ICM molecular object. The conversion algorithm adds missing heavy side-chain atoms, performs global optimization of hydrogens for best hydrogen bonding network, optimizes His, Asn, Gln, and Cys residues, and assigns partial charges. To validate the model, high affinity cPLA_2_ ligands available in ChEMBL (Target: CHEMBL3816) were docked into the binding site in the catalytic domain of cPLA_2_, and the best predicted docking scores and poses were saved for each ligand. The docking results showed reproducible binding poses and docking scores of about −25 for the known ligand scaffolds. To further improve the docking performance, the hydrogens of amino acids side chains in the binding pocket in 8 Å radius from the ligand were optimized in the presence of docked high affinity cPLA_2_ ligands. Performance of each model was evaluated based on the docking score and pose reproducibility, the value of the best docking score and an average docking score for 20 best ligands. The model optimized with molecule CHEMBL370113 showed the best reproducible results, with the best docking scores as good as - 29 and was selected for prospective screening.

#### Application of V-SYNTHES 2.0

The giga-scale screening in the REAL chemical space of 36 billion molecules was performed using the V-SYNTHES 2.0, a completely automated version of the V-SYNTHES platform described previously^11^. Docking of the pre-generated Minimal Enumeration Library (MEL), partially enumerated, and final molecules was performed with thoroughness set to 2. The docking and cheminformatic steps of V-SYNTHES 2.0 were performed in ICM-Pro (Molsoft LLC) molecular modeling software (version 3.9-2b). Docking steps were performed in the USC CARC (Center for Advanced Research Computing at the University of Southern California) facility. The cheminformatic steps of V-SYNTHES 2.0 were performed on Linux workstation (AMD Ryzen 9 7950X 16-Core Processor); 40 GB of space was needed to complete the project.

#### Selection and Synthesis of Candidate Hits for cPLA_2_

The selection of the most promising docked molecules for synthesis included the following standard post-processing steps: (i) filtering out molecules with undesirable physical-chemical properties, such as low drug-likeness, potential toxicity, and pan-assay interference (PAINS); (ii) prioritizing molecules with high predicted solubility (molLogS) and BBB penetration (iii) ensuring chemical novelty of selected molecules as compared to known high affinity ligands, using Tanimoto distance as a quantitative parameter for similarity. Also, the interactions between the predicted hits and binding pocket were analyzed and molecules with H-bond to R200, S228, T680 were given priority in the selection process. To ensure inhibition of the enzyme, molecules with close distances to catalytically active residue S228 at the bottom of the binding pocket were considered in the selection process: d < 6Å, or d = 6–8 Å with H-bonds to R200, T680, or A578. Even though the whole REAL space was screened in this work, molecules from 2-comp reactions sub-space were predicted to have the best binding due to the small and shallow nature of the cPLA2 binding pocket and were prioritized for synthesis and testing. A total of 127 compounds were selected for synthesis, out of which 117 compounds were synthesized and delivered by Enamine in 6 weeks.

#### Synthon Base Space Search for Analog Generation

The search for analogs of confirmed hits was performed in up-to-date versions of REAL Space. The analog search was performed using an in-house developed Synthon Base Space Search algorithm that takes advantage of the combinatorial nature of REAL chemical space, avoiding full enumeration of all molecules and working in the space of synthons. The algorithm starts with the decomposition of a hit molecule into the reaction scaffold and the synthons from which the compound was generated. In the next step, the algorithm searches for analogs of each of the synthons in the REAL Space using a specific Tanimoto distance (set to 0.4 in this case) as a cutoff to a molecule’s similarity. In the next step, identified synthon analogs are enumerated with each other according to the connection rules specified in REAL Space, considering physio-chemical properties to generate full molecules that comply with Lipinski’s rule of five. The Synthon Base Space Search algorithm finds tens of thousands of analogs in the Enamine REAL Space of 36 billion molecules in minutes. Generated analogs of confirmed hits from REAL Space were docked into the binding pocket of cPLA_2_, with increased effort equal to 5 to ensure comprehensive sampling of small molecules in the binding pocket. The final selection of analogs for synthesis and testing was based on the predicted docking score and the Tanimoto distance to the confirmed hit.

### cPLA2 Biochemical Testing

#### Testing using 10 ***μ***M Thresholds

Recombinant human cPLA_2_ (49 nmol, Abcam ab198469) was first pre-incubated with test compound (dissolved in DMSO) in 50 μL 1X PLA_2_ buffer (50 mM Tris-HCl, 100 mM NaCl, 1 mM CaCl_2_, pH 8.9, EnzChek™ Phospholipase A_2_ (PLA_2_) Assay Kit, Invitrogen E10217) at room temperature for 30 min. To this, 50 μL of the substrate-liposome solution [1-O-(6-BODIPY 558/568-aminohexyl)-2-BODIPY FL C5-*sn*-glycero-3-phosphocholine (Red/Green BODIPY PC-A2), dioleoylphosphatidylglycerol (DOPG), and dioleoylphosphatidylcholine (DOPC)], prepared according to manufacturer’s instructions, was added. Final concentrations were 10 μM of test compound, 600 ng/mL enzyme, and 1.6 μM fluorogenic substrate. Fluorescence (excitation at 460 nm, emission at 515 nm) was measured at 60 min from the start of the reaction and was corrected for the background fluorescence (setup with substrate but no enzyme). Enzyme activity was then calculated relative to the positive control (setup with substrate and enzyme but no inhibitor).

#### Testing using 2 ***μ***M Thresholds

Recombinant human cPLA_2_ (49 nmol, Abcam ab198469) was first pre-incubated with test compound (dissolved in DMSO) in 50 μL 1X PLA_2_ buffer (same as above) at 37°C for 40 min. To the recombinant cPLA_2_-test compound mix was added 50 μL of the substrate-liposome solution (Red/Green BODIPY PC-A2, DOPC, and DOPG), prepared according to the manufacturer’s instructions. Final concentrations were 10 μM of test compound, 800 ng/mL enzyme, and 0.55 μM fluorogenic substrate. Fluorescence (excitation at 460 nm, emission at 515 nm) was measured at 0–90 min from the start of the reaction and was corrected for background fluorescence (set up with substrate but no enzyme). The slope of the fluorescence at 30–70 min was used to calculate enzyme activity relative to the positive control (setup with substrate and enzyme but no inhibitor).

### iPLA2 Biochemical Testing

Because there is no commercial source of human recombinant calcium-independent phospholipase A_2_ (iPLA_2_), the enzyme was prepared from an overexpression cell line. Cell lysate was first pre-incubated with test compound (dissolved in DMSO) in iPLA_2_-specific buffer^12^ (100 mM Tris-HCl, pH 7.0, and 4 mM EGTA). To the recombinant iPLA_2_-test compound mix was added 50 μL of the substrate-liposome solution (Red/Green BODIPY PC-A2, DOPC, and DOPG), prepared according to the manufacturer’s instructions but replacing the buffer with 1X iPLA_2_-specific buffer. Final concentrations were 0–50 μM of test compound, 20 μg/mL lysate protein, and 0.55 μM substrate. Fluorescence (excitation at 460 nm, emission at 515 nm) was measured at 0–90 min from the start of the reaction and was corrected for background fluorescence (set up with substrate but no enzyme). The slope of the fluorescence at 30–70 min was used to calculate enzyme activity relative to the positive control (setup with substrate and enzyme but no inhibitor).

### Cell Models

Immortalized mouse astrocytes (gift from Dr. David Holtzman) were grown in DMEM/F12 (Corning, MT10090CV) containing 10 % FBS, 1 mM sodium pyruvate (Thermo Fisher, 11360070), 1 mM geneticin (Thermo Fisher, 10131-035) and 1% anti-anti. Cells were seeded in a 24-well plate (0.25×105 cells/500 μL) and cultured overnight. The cells were treated with cPLA_2_ inhibitors plus TIA (TNFα (10 ng/mL) + IFNγ (10 ng/mL) + A23187 (1 μM)) in FBS free medium for 30 minutes. After washing twice with cold PBS, cells were lysed with 1X sample buffer. Phosphorylated cPLA_2_ protein levels in cell lysate were detected by Western blot.

### In vivo Methods

#### Animals

All animal studies were approved by the Institutional Animal Care and Use Committee at the University of Southern California. The C57BL/6J mice used in the study were bred in the USC animal facility. Animals were housed with standardized 12 h light and dark cycles and had access to food and water ad libitum. Vivarium temperature was maintained between 22°C and 24°C and humidity was maintained between 50 and 60%.

#### BRI-50460 Administration and Tissue Collection

Sixteen C56BL/6 mice were randomly assigned to four groups (*n* = 4 each group) and injected with one dose of BRI-50460 (3 mg/kg, subcutaneously) formulated in an aqueous solution with 22.5% hydroxypropyl-β cyclodextrin. After the injection, animals at the designated time point (0.5, 1, 3 and 6 hours) were euthanized with isoflurane. The blood was collected into EDTA-coated tubes via cardiac puncture and plasma was isolated by centrifugation. Remaining blood was removed by transcardial perfusion with PBS, after which, the mouse brains were harvested, and flash frozen along with plasma samples for further processing.

#### BRI-50460 Quantification by LC-MS/MS

BRI-50460 was extracted from mouse plasma (100 µL) and brain tissue (150–200 mg) and quantified by liquid chromatography-tandem mass spectrometry (LC-MS/MS) using multiple reaction monitoring (MRM). Mouse brains were cut in half through the midsagittal plane prior to analysis and the left brain was used for quantitative analysis. A standard curve was prepared by spiking 3X stripped C57BL mouse plasma (BioIVT) aliquots with increasing concentrations of BRI-50460. To extract BRI-50460 and to prevent lipid oxidation, methanol (MeOH) containing 0.05% butylated hydroxytoluene and triphenylphosphine was added to plasma samples (200 µL) and brain tissues (500 µL). Each sample was also spiked with 50 µL of a structurally similar synthetic analog BRI-50469 (50 ng/mL), which was used as the internal standard. BRI-50460 was extracted from plasma samples by vortexing for 10 seconds, followed by resting on ice for 10 minutes while brain tissues were homogenized using a TissueLyser II (Qiagen) at 30 Hz for 1.5-min intervals. Subsequently, plasma and brain samples were centrifuged at 10,000×g for 10 min at 4°C, after which, supernatant was collected and diluted with water to 10% MeOH. Samples were then purified by solid-phase extraction using Strata-X 33μm Polymeric Reversed Phase cartridges (Phenomenex), dried, and prepared for LC-MS/MS analysis as previously described^13^.

BRI-50460 and the internal standard (BRI-50469) were separated and quantified in an Agilent 1290 UPLC system coupled to a Sciex API4000 system using electrospray ionization (ESI) in negative ion mode. A Poroshell 120 EC-C18 column (2.7 μm, 4.6 × 100 mm, Agilent) was used with the mobile phases: A, 0.1% formic acid in H_2_O; and B, 0.1% formic acid in MeOH. The gradient was as follows: 0–2 min, 20% B; 2–6 min, 20-95% B; 6–11 min, 95% B; 11–12 min, 95-20% B; 12–15 min, 20% B. An injection volume of 20 μL was used, the flow rate was set to 0.5 mL/min, and the column temperature was maintained at 40°C. Data analysis and quantification were performed using Skyline 23.1.3. The MRM transitions used to quantify BRI-50460, and its internal standard are detailed in **Supplementary Table S6**. The quantification of PUFAs and oxylipin metabolites was performed according to our previously described protocol^14^.

Free and protein-bound fractions of BRI-50460 were separated using Single-Use Rapid Equilibrium Dialysis Plates (Thermo Scientific) and quantified using the previously described LC-MS/MS method. To assess brain uptake, the *K*_puu_ was calculated as the ratio of unbound drug concentration in the brain divided by unbound drug concentration in the plasma^15^.

### Testing BRI-50460 in human iPSC-derived astrocytes

APOE isogenic human iPSCs were purchased from Jackson Laboratory (KOLF2.1J). Dissociated forebrain NPCs were differentiated into astrocytes in astrocyte medium (#1801, ScienCell), as previously described^16,17^. Briefly, forebrain NPCs were maintained at a high density on poly L-ornithine hydrobromide (#P3655-50MG, Sigma) and laminin (#23017015, Thermo Fisher)- coated plates and cultured in NPC medium [DMEM/F12 (Corning, MT10090CV), 1 x N2 supplement (Thermo Fisher, 17502048), 1xB27 supplement (Thermo Fisher, 12587010), 1 mg/mL laminin, and 20 ng/mL FGF2 (Thermo Fisher 13256029)]. The cells were split at approximately 1:3 to 1:4 every week with Accutase (Sigma, SCR005). NPCs were differentiated into astrocytes by seeding dissociated single cells at a density of 15,000 cells/cm^2^ on Matrigel (Corning, 356255) -coated plates and cultured in complete astrocyte medium (#1801, ScienCell). After 30 days of differentiation, the astrocytes were ready for further experiments. Mature APOE ε3/ε3 astrocytes were seeded at 0.5×10^5^ cells/well in a 24-well plate and grown to 70-80% confluency. Cells were washed with FBS-free medium and treated with or without 0.1 or 1µM BRI-50460 for 30 minutes. Then, cells were treated with or without 2.5 µM amyloid beta 1-42 (Anaspec) for 72 hours. After treatment, cells were lysed with radioimmunoprecipitation assay (RIPA) buffer (9806, Cell Signaling Technology, CST), and total cPLA_2_ and phosphorylated cPLA_2_ protein levels were detected by western blotting.

### Testing BRI-50460 in human induced pluripotent stem cells (iPSCs) and Neural progenitor cells (NPCs)-derived AD cellular models

Neural progenitor cells (NPCs) were differentiated from iPSCs using the StemCell Technologies STEMdiff™ SMADi Neural Induction Kit (Catalog # 08581) following manufacturer’s protocol. Briefly, JAX IPSC 1162 (APOE3/3) and JAX IPSC 1150 (APOE4/4) iPSCs were grown on Matrigel (Corning Catalog # 354230)-coated dishes in mTeSR™ Plus (StemCell Technologies Catalog # 100-0276). To initiate NPC induction, iPSCs were passaged to either Matrigel- or Poly-L-Ornithine hydrobromide (PORN; Sigma Aldrich P3655)/laminin (ThermoFisher 23017015)-coated plates and were fed with STEMdiff™ Neural Induction Medium + SMADi. At passage 3, cells were switched to STEMdiff™ Neural Progenitor Medium (StemCell Technologies Catalog #05833) and were maintained in that medium. For neuronal differentiation, NPCs were seeded on PORN/laminin-coated chamber slides at 20,0000 cells/cm^2^. For neuronal induction, cells were fed with the neuronal medium as described in Bardy et al.^18^ At four to five weeks of neuronal differentiation, the cells were treated with 2.5 µM amyloid-beta 42 oligomers with or without BRI-50460 for 72 hours, then the cells were fixed in 4% PFA for 15 mins and stained with antibodies: phospho-cPLA2Ser505 (1:100, Sigma Aldrich Catalog # SAB4503812), phospho-tau Ser202/Thr205 (AT8) (1: 300, ThermoFisher Catalog # MN1020), CaMKIIα (1: 300, Catalog # 50049, Cell Signaling), synapsin I (1: 300, Sigma Aldrich Catalog # AB1543), and MAP2 (1: 1000, ThermoFisher Catalog # PA1-10005). All images were taken under Leica TCS SP8 confocal microscope. The images were processed and analyzed using ImageJ and CellProfiler.

## Results

### Giga-Scale virtual screen for BBB penetrant cPLA2 inhibitors with V-SYNTHES 2.0

We used the V-SYNTHES 2.0 platform to virtually screen the Enamine REAL chemical space to identify potential small-molecule inhibitors of cPLA_2_. Application of the V-SYNTHES 2.0 enabled effective screening of the 36-billion compound chemical space, while performing the docking of only a small fraction of the molecules (∼2 million). The algorithm starts with the docking of a pre-generated Minimal Enumeration Library (MEL) to the cPLA_2_ binding pocket. MEL represents the whole chemical space, with ∼1 million fragment-like combinations of every reaction scaffold with one real synthon from the chemical space, and minimal “caps” that substitute the real counter-synthons. The best MEL compounds are selected based on the cPLA_2_ docking score and molecule pose in the binding pocket and enumerated with the second real synthon. For 2-component reaction sub-space, full molecules are generated after one round of enumeration: for 3-component reactions sub-space – after two rounds of optimization. The full molecules are then docked into the cPLA_2_ binding pocket, and the best scoring candidates are analyzed to make the final selection of compounds for synthesis and testing.

A giga-scale virtual screening was performed using a ligand-optimized cPLA_2_ structural model based on the apo-enzyme structure (PDB ID 1CJY, **Fig. 1**). Candidate hit compounds were selected based on predicted docking scores and binding poses, chemical novelty, and diversity, and filtered for low PAIN and toxicity scores, as well as high blood-brain barrier (BBB) penetration properties. A total of 127 top-scoring compounds (**Supplementary Table S1**) were selected for synthesis, out of which 117 compounds were synthesized by Enamine in less than 6 weeks.

**Figure 1:**
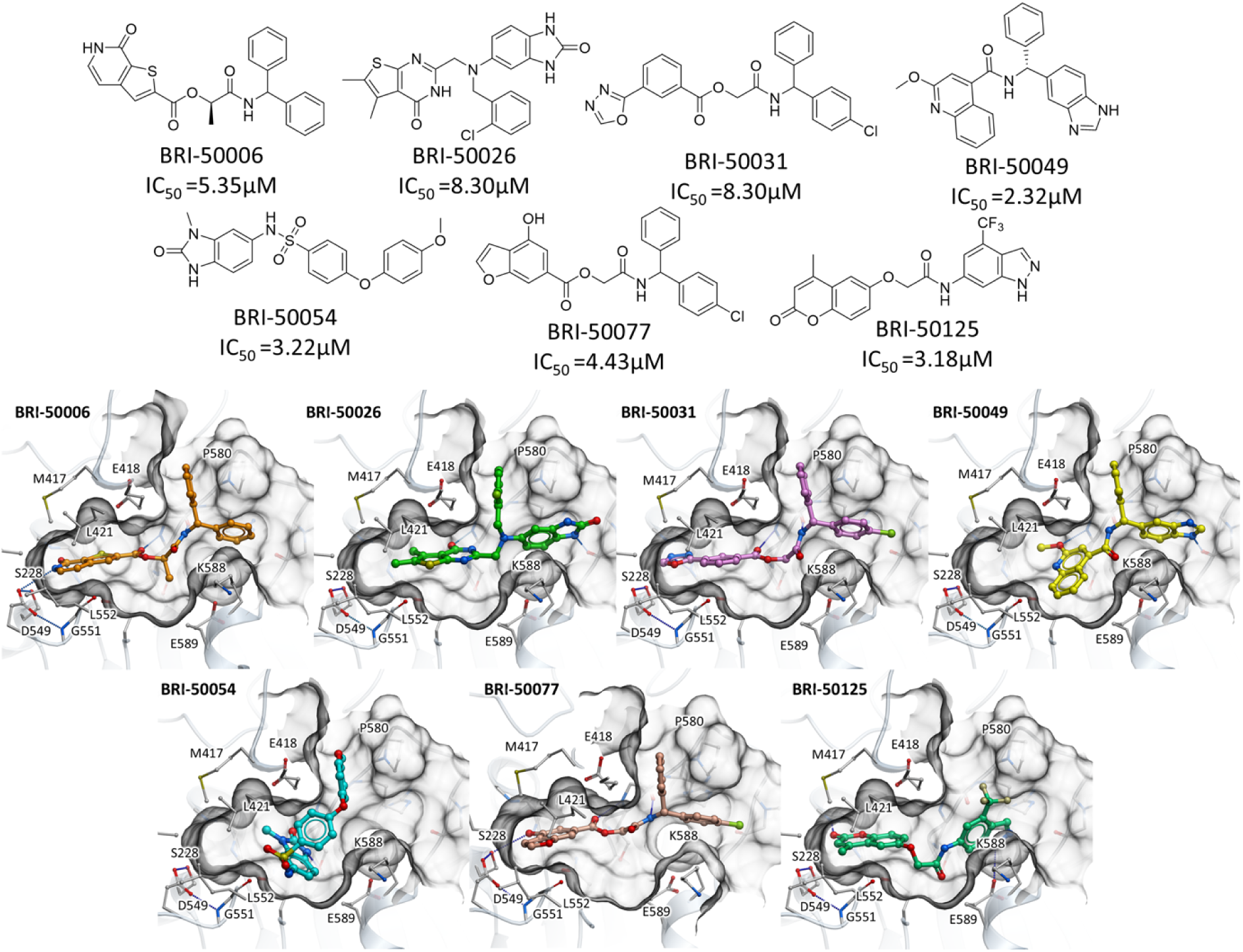
Predicted binding poses of compounds on the cPLA_2_ crystal structure (PDB ID 1CJY) from the 1st Generation selected for potency screening optimization for cPLA_2_ inhibition.

### Identification of the Initial High-quality cPLA_2_ Inhibitor Hits

To test the predicted hit candidates in vitro, we used a cPLA_2_ inhibition fluorescence assay (**Fig. 2A**), with commercially available cPLA_2_ inhibitor ASB14780 serving as the positive control. This initial screen identified 19 compounds with >40% inhibition of cPLA_2_ at 10 μM for a prospective screening rate of 16%. Dose-response curves measured for the best 7 new compound chemotypes (**Supplementary Table S2**) showed IC_50_ values below 10 μM, comparable to our reference inhibitor ASB14780 with a determined IC_50_ = 6.66 μM in the same assay.

**Figure 2:**
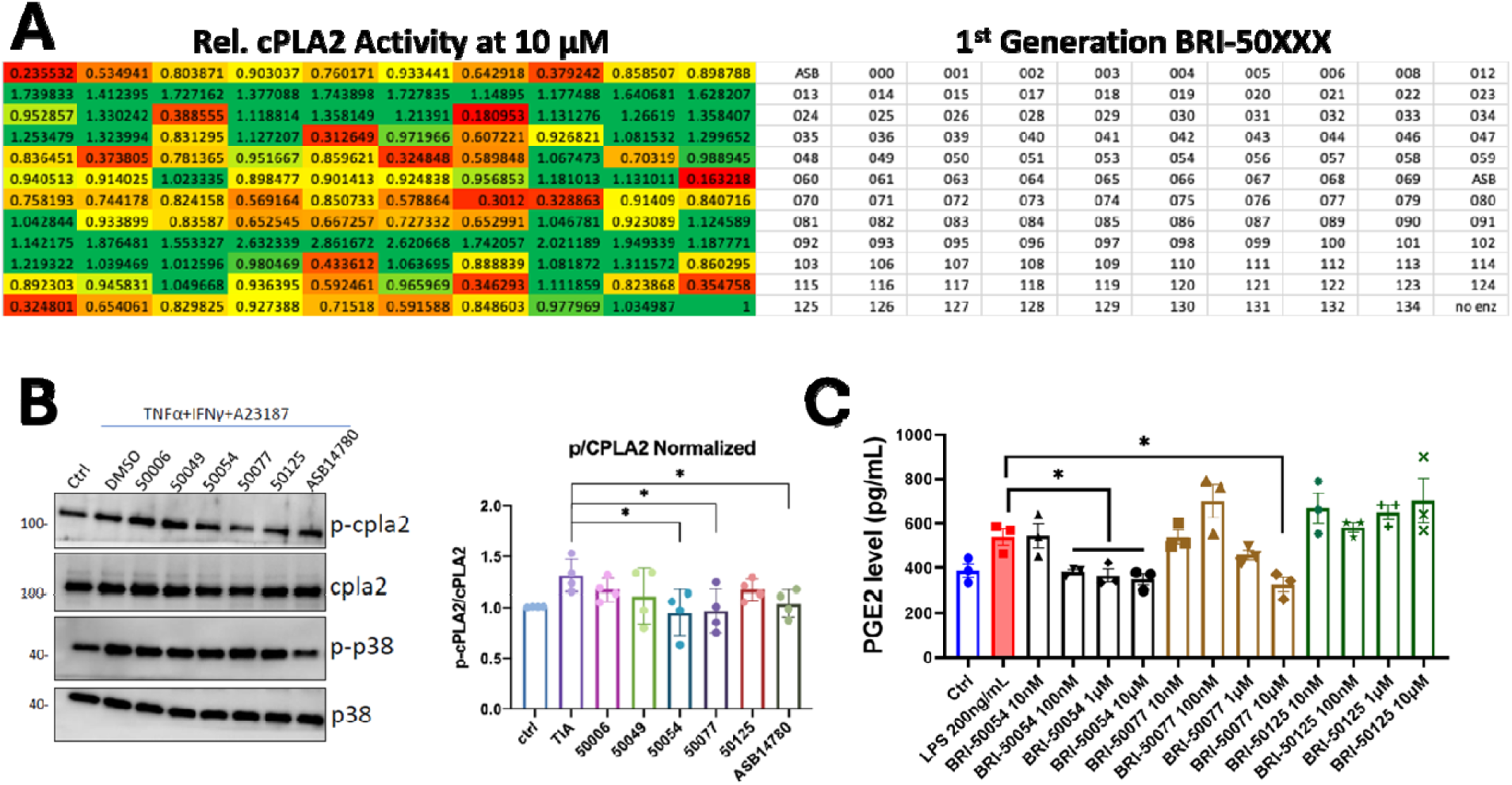
Preliminary cPLA_2_ inhibition screen of 1^st^ Generation virtual screening hits. (**A**) A fluorescence-based assay was used to determine the activity of recombinant human cPLA_2_ incubated with 10 μM test compound relative to a control (substrate, enzyme, no compound) in our initial screen. ASB14780 was also included as an inhibitor positive control. Compounds BRI-50006, BRI-50049, BRI-50054, BRI-50077, and BRI-50125 were among the hits found through this initial screen. **(B)** The compounds (10 µM) were mixed with stimuli (TIA: TNFa+rIFN+A23187) and added to astrocytes for 30 minutes. The cell lysate was harvested and used to test the protein levels of phosphorylated and total cPLA_s_ by Western blot. ASB14780 was used as a positive cPLA_2_ inhibitor control. (**C**) BV-2 microglia cells were pre-treated with different concentration of BRI-50054, BRI-50077 and BRI-50125 for 30 minutes, then treated with LPS (10 ng/mL) for 16 hours. PGE_2_ levels in the medium was measured using a PGE_2_ ELISA assay kit.

Confirmed cPLA_2_ inhibitors have diverse chemical scaffolds with a Tanimoto distance >0.3 between each other and >0.4 to known cPLA_2_ inhibitors. **Figure 1** shows the diversity of predicted docking poses for the 7 hit compounds. Compounds BRI-50006, BRI-50031 and BRI-50077 have similar moieties on one side of the molecule; however, the second synthon adds diversity to the chemical scaffold and ensures a variety of interactions between the ligand and the residues of the binding pocket. In all three molecules, carbonyl oxygen from the ether group and nitrogen from the amide group form hydrogen bond interactions with the backbone of catalytically active residue A578. The other parts of the 3 hit molecules cover different interactions: for instance, compound BRI-50006 forms hydrogen bonds with residue D549, BRI-50031 bonds with the backbone of residue G197, and BRI-50077 with residue S228. In a similar manner, the carbonyl oxygen of BRI-50054 and the methoxy group of BRI-50049 form hydrogen bonds with the backbone of residue A578. BRI-50125 forms hydrogen bonds with residue E589 and the backbone of G197.

Subsequent cellular screening revealed that BRI-50054 showed potent inhibition and reduced p-cPLA_2_/cPLA_2_ levels (**Fig. 2B**) as well as reduced downstream production of PGE_2_ in cell models of neuroinflammation (**Fig. 2C**), with minimal impact on mitogen-activated protein kinase (MAPK) pathways. These *in vitro* hits were further assessed using animal models, where the pharmacokinetics of each compound were evaluated *in vivo* for their bioavailability, drug distribution in various organs, metabolic stability, and various PK parameters. Compound plasma and brain concentrations were measured at multiple timepoints. This integrative approach provided the basis for selecting compounds for further development and testing in animal models.

Furthermore, pharmacokinetic studies in C57BL/6 mice revealed that the 1^st^ Generation compound BRI-50054 was well-tolerated and exhibited dose-dependent increases in both the plasma and brain. This was accompanied by reduced brain AA levels and elevated levels of ω-3 fatty acids (DHA and EPA, **Fig. 3B, D**) and their pro-resolving metabolites, such as resolvins.

**Figure 3:**
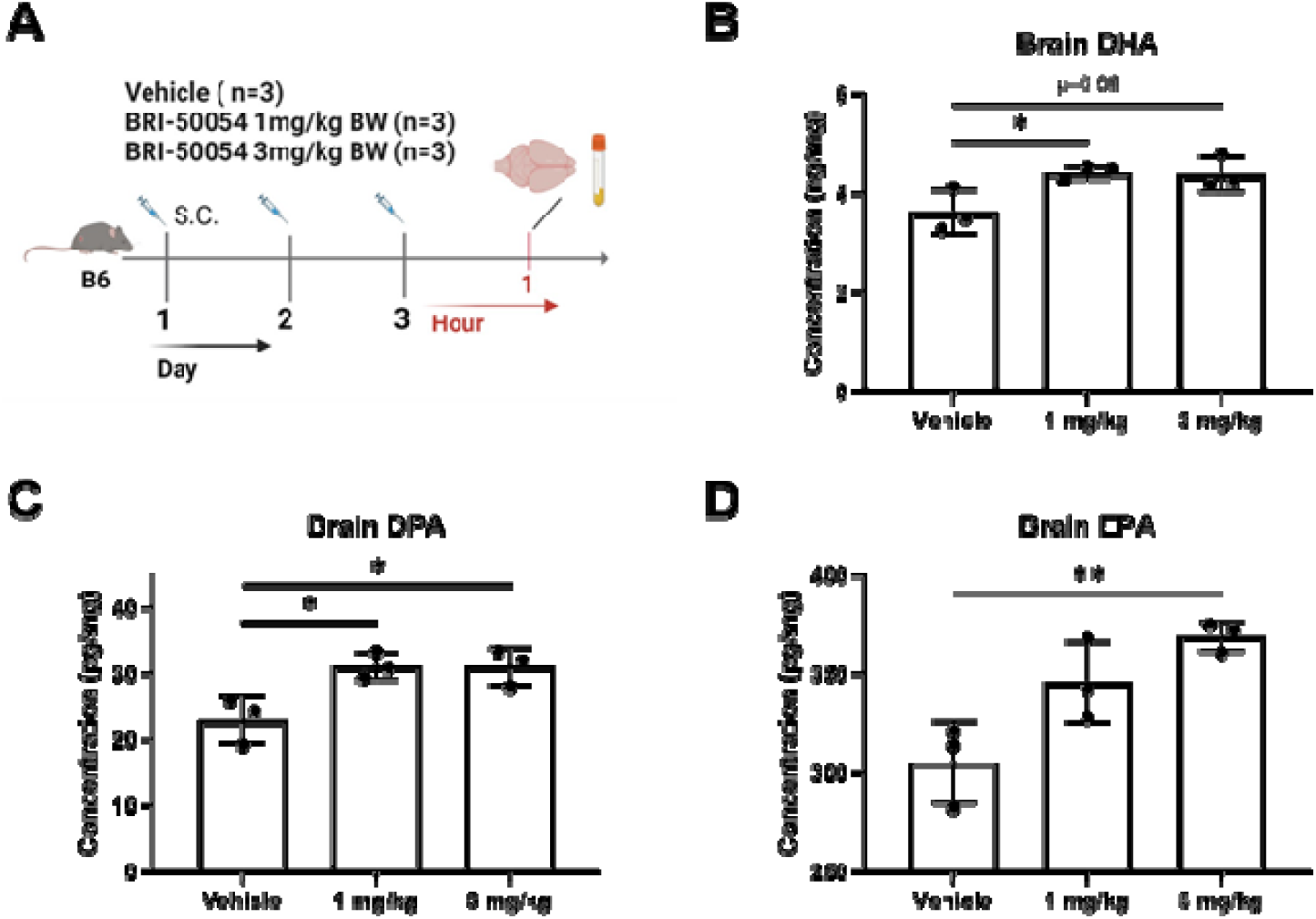
BRI-50054 Treatment Increases Free ω-3 PUFA Brain Levels. (**A**) C57BL/6J mice (n=3) treated with vehicle, 1.0 or 3.0 mg/kg/day IP BRI-50054 for 3 days. Brain levels of DHA (**B**), DPA (**C**) and EPA (**D**) were measured using a validated LC-MS/MS lipidomic assay. * p<0.05, ** p< 0.01, *** p < 0.001

These findings support the ability of cPLA_2_ inhibitors to engage their target *in vivo*, restore brain lipid homeostasis, and mitigate the neuroinflammatory environment characteristic of Alzheimer’s disease (AD).

### SAR-by-Catalog and the First Round of Hit Optimization

Hit compounds identified using the V-SYNTHES 2.0 algorithm can be effectively optimized using on-demand synthesis, as the combinatorial nature of REAL space contains thousands of readily available close analogs. Based on the *in vitro* and *in vivo* testing results, molecules BRI-50054, BRI-50077 and BRI-50125 were nominated as the most promising hits for optimization. The structure-activity relationships (SAR)-by-catalog was performed using an in-house developed Synthon Base Space Search algorithm that emerged as a stand-alone tool for chemical search in REAL space. Like V-SYNTHES2.0, this search algorithm effectively leverages the combinatorial nature of REAL Space. The search algorithm decomposes hit compounds into a reaction scaffold and the synthons from which the compound was generated (**Supplementary Fig. S1**) and performs a search of synthon analogs in the REAL space. Then, full analog molecules are generated by enumeration of identified synthon analogs according to the connection rules specified in REAL Space. This synthon search algorithm avoids full enumeration of the chemical space and takes only minutes to find tens of thousands of hit analogs among >36 billion REAL Space compounds. Based on docking scores and predicted physio-chemical properties like solubility, PAIN, and BBB penetration, a total of 100 analogs for hit compounds BRI-50054, BRI-50077, and BRI-50125 were selected for synthesis (**Supplementary Table S3**). Out of 100 requested, 85 molecules were synthesized by Enamine in <6 weeks and experimentally tested.

Using a cellular assay in astrocytes stimulated with TNFα + IFNγ + A23187 (TIA) to induce cPLA2 phosphorylation, inhibitory effects of these 2^nd^ Generation analogs were assessed via Western blot (**Supplementary Fig. S2A**). From the initial screen at 10 μM, 39 compounds were selected for further evaluation at 5 μM (**Supplementary Fig. S2B**). Subsequently, 10 compounds were tested at 1 μM (**Supplementary Fig. S2C**), identifying four most promising candidates: BRI-50202, BRI-50206, BRI-50229, and BRI-50286. These compounds demonstrated significant inhibition of cPLA_2_ activation, validating their potential as optimized leads.

### Second Round: Lead Optimization

Based on the results from the initial screening and SAR-by-catalog, two scaffolds were selected for further optimization: first represented by BRI-50054 and BRI-50202 compounds, and second – by BRI-50125 and BRI-50281 compounds. With the priority given to the scaffold represented by hits BRI-50054 and BRI-50202 (**Fig. 4C**), the search for analogs was performed in REAL space of 36 billion molecules using the previously described developed in-house chemical search engine. After analysis of docking results for generated analogs, 113 analogs were requested for synthesis from Enamine. 103 out of 113 requested compounds (**Supplementary Table S4**) were synthetized in 6 weeks and experimentally tested.

**Figure 4:**
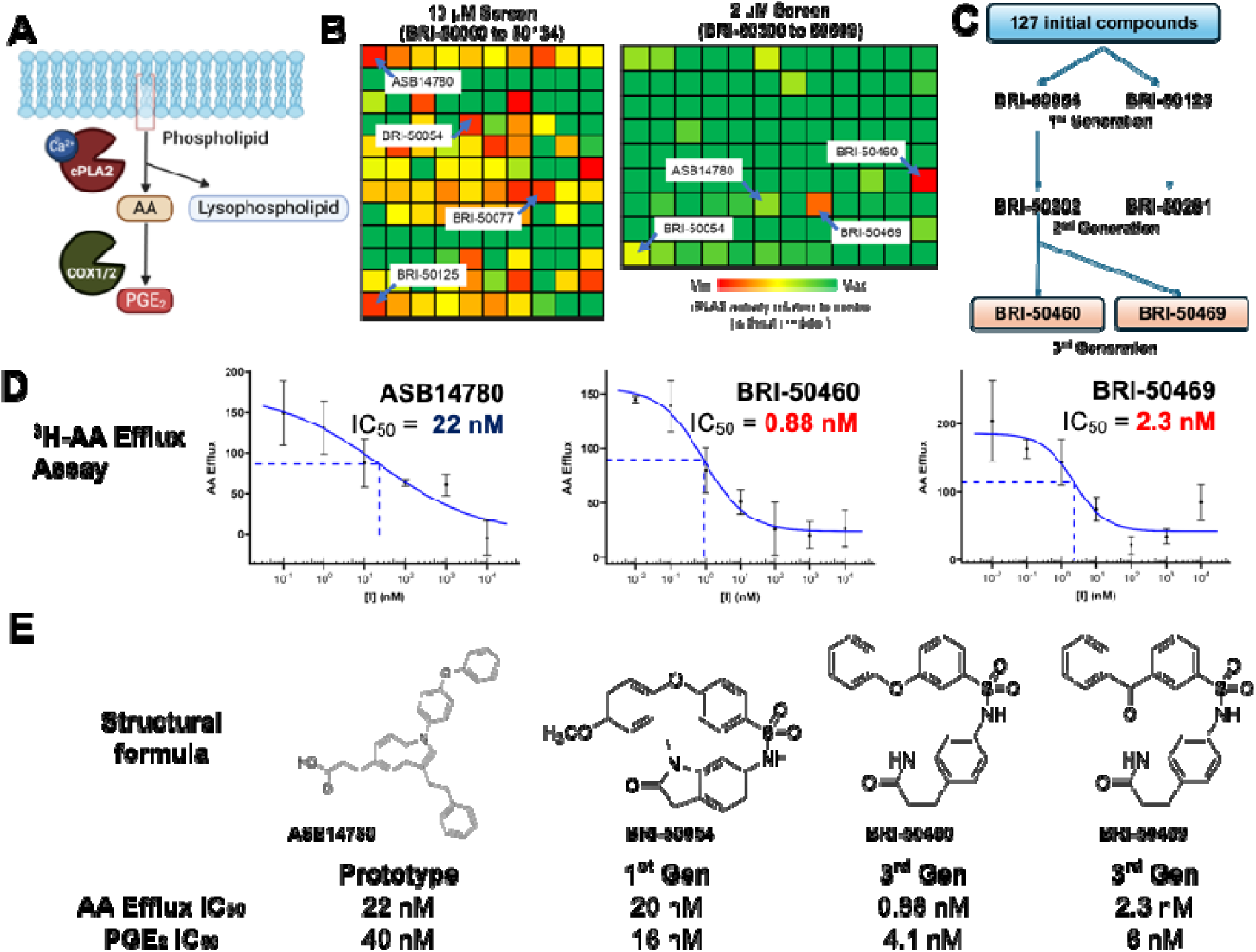
*In vitro* Inhibition Data. (**A**) Schematic for the reaction of cPLA2 on membrane phospholipid to release AA. The released AA is converted to PGE_2_ by COX enzymes. (**B**) Preliminary cPLA2 in vitro activity screens of the first and third generation of small molecules. Each square is the cPLA2 in the presence of inhibitor relative to control (no inhibitor). (**C**) Progression of iterative structure optimization and potency screening of cPLA2 inhibitors. (**D**) Dose-dependency curve for the inhibition of AA efflux by ASB14780 and our two 3^rd^ Gen leads upon administration of A23187 (Ca^2+^ ionophore). (**E**) IC_50_ estimates for the inhibition of AA efflux into cells and the inhibition of PGE_2_ production.

The second round of SAR-by-catalog and in vitro screening with a 2 μM threshold (**Fig. 4B**) yielded 3^rd^ Generation compounds BRI-50460 and BRI-50469, which exhibited superior *in vitro* potency compared to earlier candidates. Both compounds demonstrated high efficacy in inhibiting AA release from cells labeled with ^3^H-AA and treated with a calcium ionophore (A23187). Among them, BRI-50460 emerged as the most potent inhibitor, with an IC_50_ of 0.88 nM for cellular AA release and an IC_50_ of 4.1 nM for downstream PGE_2_ inhibition, representing a >20-fold improvement over ASB14780 (**Figs. 4D, E, Supplementary Table S5**). We also observed high selectivity for cPLA_2_ over iPLA_2_ in our *in vitro* assays; at 50 μM or either BRI-50460 or BRI-50460, iPLA_2_ was reduced by only *ca.* 23% (**Fig. 5**).

**Figure 5:**
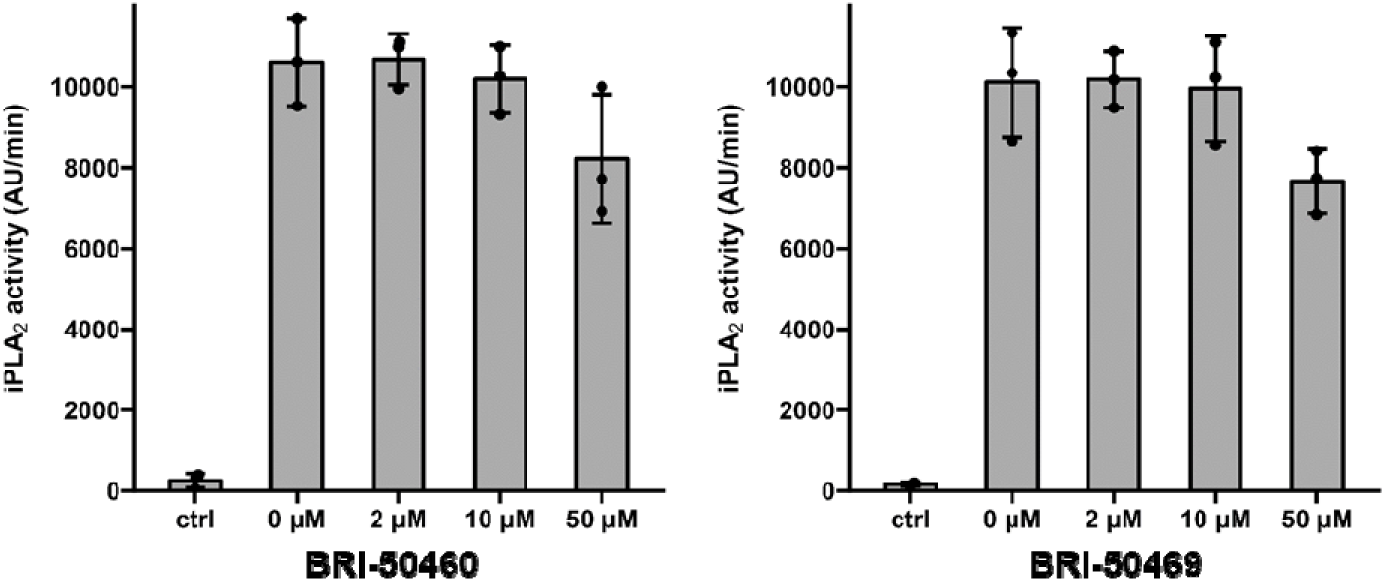
In vitro iPLA_2_ inhibition assay on two lead compounds BRI-50460 and BRI-50469.

### Brain Delivery and Pharmacokinetics of BRI-50460

In an acute pharmacokinetic study, BRI-50460 was administered subcutaneously to C57BL/6 mice (3 mg/kg body weight). Plasma and brain tissue analysis revealed a brain-to-plasma (B/P) ratio exceeding 40%, more than double that of the 1^st^ Generation compound BRI-50054 (**Fig. 6B**). Free fractions of BRI-50460 in the brain (3.28%) and plasma (0.39%) yielded a calculated *K*_p,uu_ of 2.94, indicative of robust central nervous system (CNS) penetration. This brain penetration surpassed many known CNS-penetrating drugs, including Zolpidem (Ambien) (**Fig. 6D**). However, the compound exhibited a moderate half-life, warranting further optimization for sustained therapeutic efficacy.

**Figure 6:**
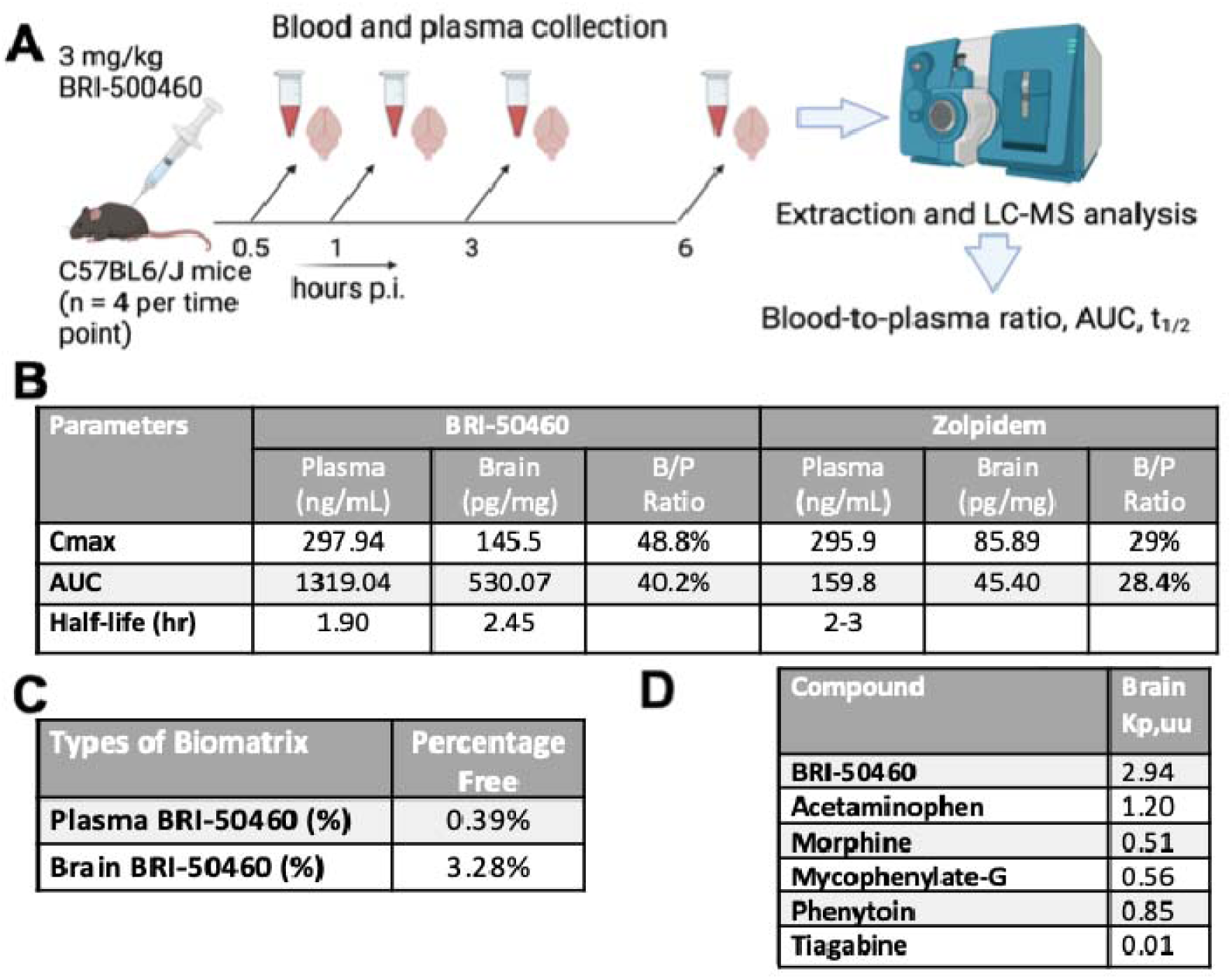
BRI-50460 Plasma and Brain Distribution. (**A**) BRI-50460 (3 mg/kg SQ) in an acute PK study using C57BL/6J mice (*n* = 4 per time point), where plasma and brain tissue were harvested at 0.5, 1, 3, and 6 hours post-injection (**B**) Plasma and brain Cmax and AUC after 3 mg/kg SQ BRI-50460 with Brain/plasma (B/P) ratio compared with zolpidem (Ambien®) (**C**) Free plasma and brain levels of BRI-50460 expressed in percentage unbound. (**D**) *K*_p,uu_ of BRI-50460 compared to other CNS penetrating agents.

### BRI-50460 reduces AB42-induced cPLA_2_ levels and cPLA_2_ phosphorylation in human iPSC-derived astrocytes

To assess whether BRI-50460 has an inhibitory effect on cPLA_2_ phosphorylation induced by AD pathology *in vitro*, human iPSC-derived astrocytes were first treated with 0.1 or 1 µM of BRI-50460 for 30 minutes, followed by 2.5 µM of Aβ42O for 72 hours. Results show that 1 μM of BRI-50460 can reduce amyloid beta (Aβ)-related increases in cPLA_2_ levels (**Fig. 7**), as well as cause a reduction in phosphorylated cPLA_2_. These data indicate that in parallel with our initial *in vitro* screening performed in immortalized mouse astrocytes, BRI-50460 also has an inhibitory effect on cPLA_2_ and phosphorylated cPLA_2_ in human iPSC-derived astrocytes challenged with Aβ.

**Figure 7:**
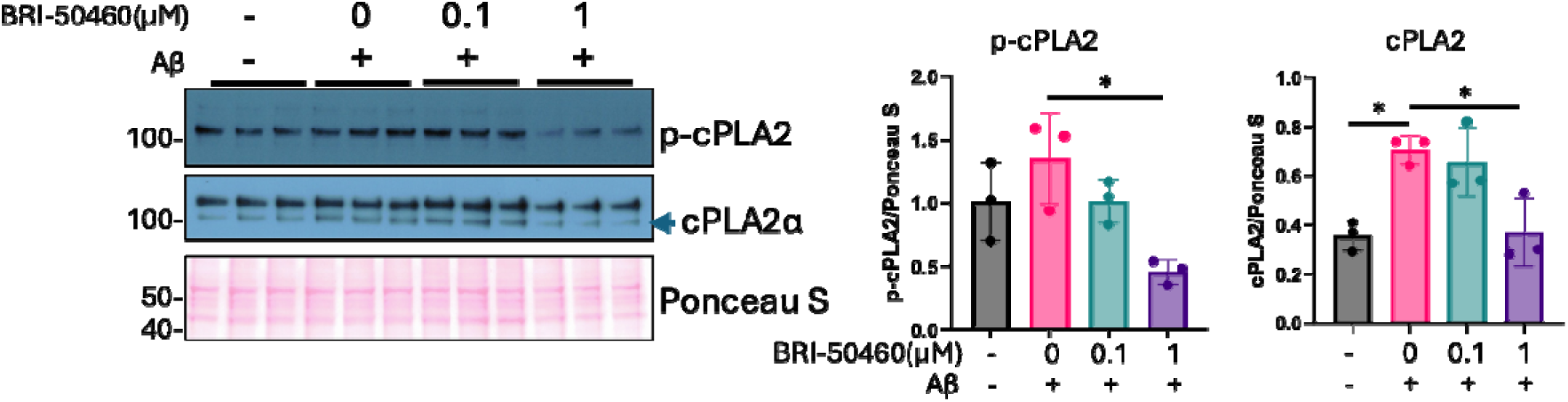
At 1μM, BRI-50460 reduces amyloid-beta42-induced cPLA_2_ levels and lowers cPLA_2_ phosphorylation in iPSC-derived astrocytes. Phosphorylated cPLA_2_ and total cPLA_2_ (indicated by arrow) levels in cell extracts from iPSC-derived astrocytes were detected by Western blot. Quantification of total cPLA_2_ and p-cPLA_2_ protein levels normalized to Ponceau stain. Data are represented as mean ± SD and analyzed using one-way ANOVA followed by Tukey’s test. * p < 0.05, **p < 0.01, ****p < 0.0001

### BRI-50460 mitigates the effects of amyloid-42 oligomers on cPLA_2_ activation, tau hyperphosphorylation, and synaptic loss in neurons derived from human iPSCs/NPCs

To examine BRI-50460’s treatment effects on AD pathologies, we first examined its impact on Aβ42O-induced tau hyperphosphorylation in a mixed-culture of neurons and astrocytes derived from human iPSCs carrying *APOE3/3* genotype. Abnormal hyperphosphorylated tau leads to the formation of neurofibrillary tangles and neurodegeneration, a pathological feature of AD^19^. After neural progenitor cells (NPCs) were differentiated for five weeks^18^, the cells were exposed to 2.5 µM of Aβ42O, with or without 1 µM BRI-50460 for 72 hours. Immunofluorescence staining revealed that BRI-50460 reduced Aβ42O-induced increases in phospho-cPLA_2_α(Ser505) (p-cPLA_2_α, **Fig. 8A, B**) and phospho-tau (Ser202/Thr205) (p-Tau, **Fig. 8A, C**) in human neurons.

**Figure 8:**
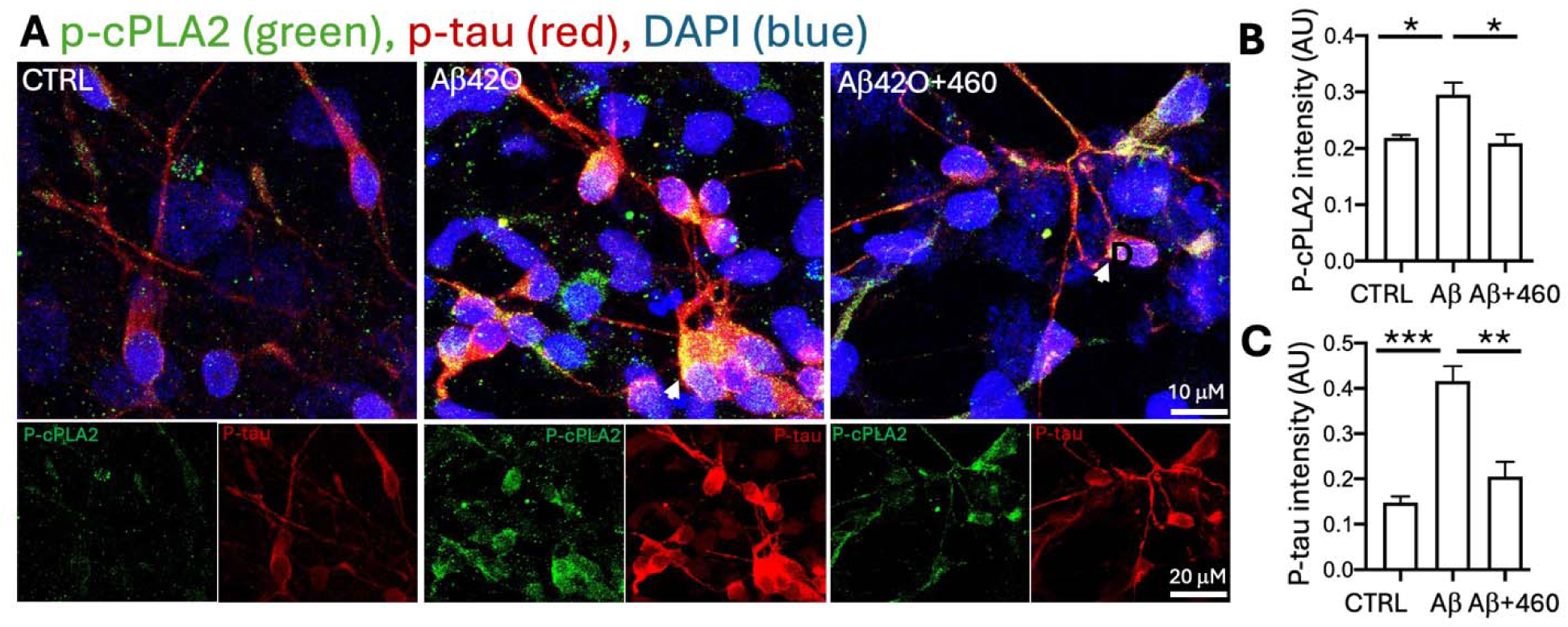
BRI-50460 amyloid-42 oligomer-induced cPLA2α activation and tau phosphorylation in neurons derived from human iPSCs. **A.** Representative images of the immunofluorescent staining of p-cPLA_2_α and p-Tau in human iPSC-derived neurons treated with 2.5 μM Aβ42 oligomers (Aβ42O) with or without 1 μM BRI-50460 for 72 hours. Quantification revealed that BRI-50460 (460) reduced Aβ42O-induced increases in p-cPLA_2_α (**B**) and p-tau (**C**) intensity. * p < 0.05, ** p < 0.01, *** p < 0.001, **** p < 0.0001. Analyzed image numbers include CTRL (*n* = 13), Aβ42O (*n* = 14), 460 (*n* = 14).

### BRI-50460 Protects Synaptic Loss in Amyloid Beta 42 Oligomer-challenged Neurons Derived from Human iPSCs

Since synaptic loss predicts early memory impairment and correlates with cognitive decline in AD^20^ and is associated with neuroinflammation^21^, we next examined whether BRI-50460 has treatment effects on Aβ42O-induced synaptic loss in the mixed-culture of neurons and astrocytes derived from human iPSCs carrying *APOE4/4* genotype. After neural progenitor cells (NPCs) were differentiated for four weeks, the cells were exposed to 2.5 µM of Aβ42O, with or without 1 µM BRI-50460 for 72 hours. Immunofluorescence staining revealed that BRI-50460 protected the cells from Aβ42O-induced reductions of synaptic puncta stained by presynaptic protein synapsin (**Fig. 9A, B**), and dendritic density (occupied area) stained by postsynaptic protein CamKIIα (**Fig. 9A, C**) and MAP2 (**Fig. 9A, D**). Synapsin regulates neurotransmitters release at synapses^22^. CaMKIIa regulates long-term potentiation, a cellular mechanism of synaptic strength underlying learning and memory^23–25^.

**Figure 9:**
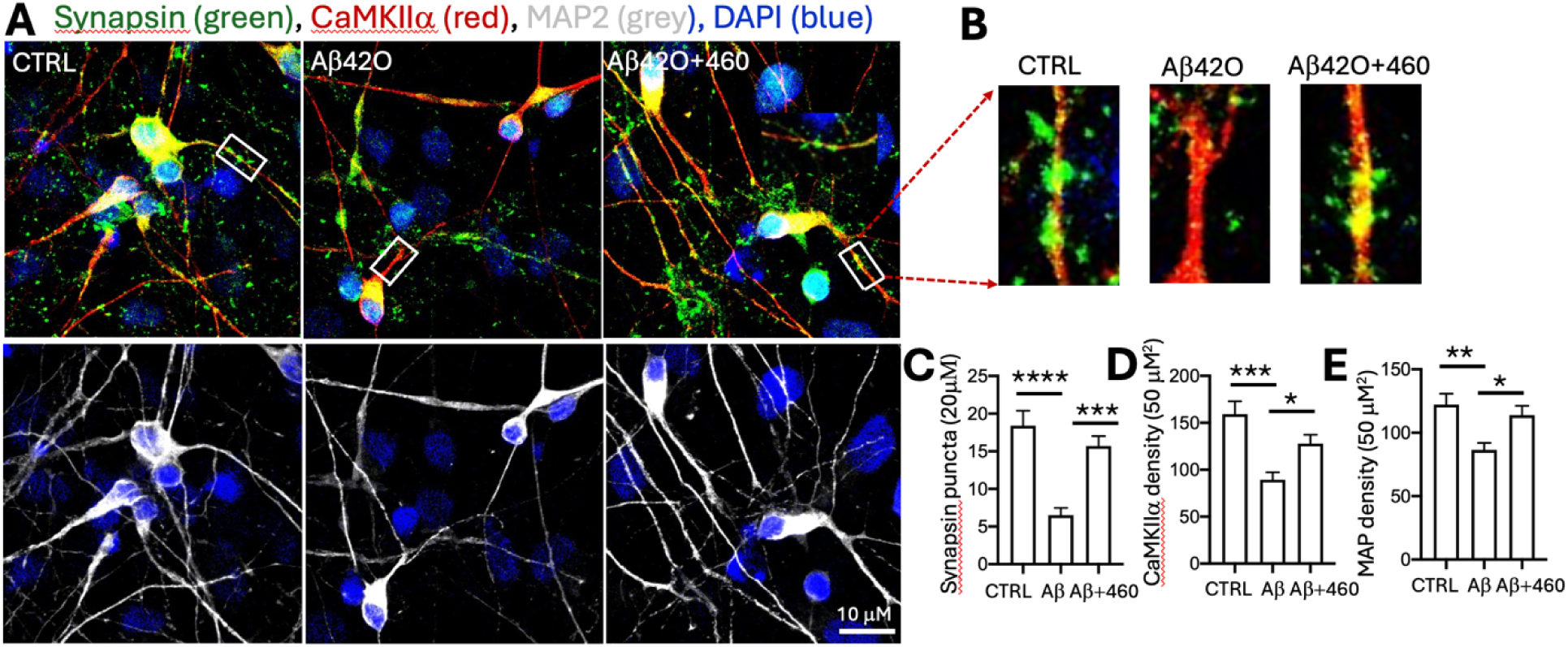
BRI-50460 protected amyloid-42 oligomer-synaptic and dendritic reduction in neuron derived from human iPSCs. (**A**) Representative images of the immunofluorescent staining of synapsin (green), CamKII (red), and MAP2 (white) in iPSC-derived neurons treated with 2.5 μM Aβ42 oligomer (Aβ42O) with or without 1μM BRI-50460 for 72 hours. (**B)** Enlarged images of the boxed dendrites in A. BRI-50460 (460) reduced Aβ42O-induced reduction of synapsin puncta (**C**), dendritic CamKIIa (**D**) and MAP2 (**E**) density in neurons derived from human iPSCs. * p < 0.05, ** p < 0.01, *** p < 0.001, **** p < 0.0001. Analyzed image numbers for CamKIIα and MAP2 include CTRL (*n* = 12), Aβ42O (*n* = 13), 460 (*n* = 13), for synapsin puncta include CTRL (194), Aβ42O (*n* = 68), 460 (*n* = 167).

## Discussion

Our structure-based virtual screening and SAR-by catalog optimization efforts have identified BRI-50460 as a highly promising cPLA_2_ inhibitor, demonstrating significant potential as therapeutic candidate for addressing neuroinflammation and pathology of Alzheimer’s disease (AD). Among the compounds derived from our structure-based cPLA_2_ virtual screening, BRI-50460 exhibits high potency, a brain penetrant pharmacokinetic profile, and an ability to modulate key mediators involved in neuroinflammation and AD pathology *in vitro*.

BRI-50460 exhibited exceptional potency, with an IC_50_ of 0.88 nM for inhibiting AA release and of 4.1 nM for blocking downstream PGE_2_ synthesis, representing a greater than 20-fold improvement over ASB14780, a previously studied cPLA_2_ inhibitor, while also demonstrating high selectivity over iPLA_2_. This enhanced efficacy underscores the success of our optimization strategy, which focused on improving both binding affinity and functional inhibition of cPLA_2_ in relevant cellular models. BRI-50460 demonstrated robust inhibition in cell-based assays without significantly affecting MAPK pathways, suggesting a targeted action on cPLA_2_ without disrupting other critical signaling cascades, which could be important for minimizing off-target effects in clinical applications.

In addition to its potent inhibition of cPLA_2_, BRI-50460 showed promising pharmacokinetic properties that are critical for CNS drug development. Upon subcutaneous administration to C57BL/6J mice, BRI-50460 achieved a brain-to-plasma (B/P) ratio exceeding 40%, more than double that of the first-generation compound BRI-50054. This indicates that BRI-50460 has superior brain penetration, a crucial feature for targeting neuroinflammation in AD. Furthermore, BRI-50460 exhibited a free fraction of 3.28% in the brain and 0.39% in plasma, yielding a calculated *K*_p,uu_ of 2.94, which further supports its potential for effective central nervous system (CNS) targeting. These values surpass those of several known CNS-penetrating drugs, including zolpidem, highlighting BRI-50460’s promising CNS bioavailability.

The ability of the 1^st^ Generation compound BRI-50054 to reduce brain AA levels and increase ω-3 fatty acids (DHA and EPA), as well as their pro-resolving metabolites like resolvins, is particularly noteworthy. This shift in lipid homeostasis could help mitigate the neuroinflammatory environment characteristic of AD, offering a therapeutic strategy that not only targets the enzyme itself but also modulates the downstream lipid signaling pathways associated with AD pathology. These findings provide strong evidence that BRI-50460 may also effectively engage its target *in vivo*, restore lipid homeostasis in the brain, and potentially alleviate neuroinflammation—a key hallmark of Alzheimer’s disease. Additionally, BRI-50460 counteracts the impact of Aβ42O on cPLA_2_ activation, tau phosphorylation, and synaptic loss in astrocytes and neurons derived from human iPSCs. This is especially interesting for restoring CaMKIIα, an essential postsynaptic protein that regulates learning and memory^23,25^.

The enzyme cPLA_2_ is notable for releasing AA, which plays an important role in neurotransmission and serves as a precursor for eicosanoids^26,27^. However, excessive activation of cPLA_2_ can accelerate the cyclooxygenase (COX) and lipoxygenase (LOX)-mediated metabolism of free AA to the products prostaglandins (PGs) and leukotrienes (LTs), respectively, which are potent inflammatory lipid mediators^6,28^. The activity of cPLA_2_ is regulated by phosphorylation and is influenced by the MAPK, particularly through phosphorylated p38^29,30^.

We have identified the activation of cPLA_2_ as a critical driver of neuroinflammation in AD, particularly in APOE4 carriers^7^. This highlights the therapeutic potential of targeting cPLA_2_ to mitigate the harmful effects of chronic inflammation in the AD brain. Other groups have reported elevated levels of cPLA_2_ protein and its phosphorylated form in astrocytes surrounding Aβ plaques compared to healthy controls^31–33^. Increased activation of cPLA_2_ is also observed in the hippocampus of human amyloid precursor protein transgenic mice^32^. In addition, Aβ oligomers can activate cPLA_2_, thereby promoting neurodegeneration^34,35^. Importantly, the Mucke group reported that cPLA_2_ genetic deficiency ameliorates memory impairment and hyperactivated glial cells observed in an AD mouse model (**Fig. 2**)^32,36^. Knock-out of cPLA_2_ in microglia can decrease lipopolysaccharide-induced inflammatory response^37^. In APOE-targeted replacement mice and human brains, we observed that APOE4 induces greater MAPK p38-mediated activation of cPLA_2_. Importantly, inhibiting cPLA_2_ reduced the release of inflammatory neurotoxic lipids and eicosanoids^7^, corresponding to reduction in neuroinflammation and synaptic loss.

## Conclusion

Our findings reveal a successful application of fast structure-based giga-scale V-SYNTHES 2.0 screening and optimization to discover new potent cPLA_2_ inhibitor chemotypes. This includes BRI-50460, which exhibits enhanced potency, selectivity, pharmacodynamic, and pharmacokinetic profiles compared to the previously developed inhibitor ASB14780. BRI-50460 demonstrated strong *in vitro* inhibition of cPLA_2_-mediated arachidonic acid (AA) release, with sub-nanomolar IC_50_ value and improved blood-brain barrier (BBB) penetration measures. By inhibiting cPLA_2_, BRI-50460 has the potential to restore lipid homeostasis, reduce AD pathology and synaptic dysfunction, particularly in APOE4 models, making it a promising candidate for further clinical development.

This study underscores the potential of advanced virtual screening platforms like V-SYNTHES 2.0 in accelerating the discovery of CNS-targeting therapeutics and lays the groundwork for further exploration of BRI-50460 in chronic treatment paradigms with a prominent cPLA_2_ activation profile. Ongoing efforts will focus on refining its pharmacokinetics and evaluating its therapeutic effects in preclinical AD models, with the goal of advancing it toward clinical evaluation for the treatment of neuroinflammatory conditions in Alzheimer’s disease.

## Abbreviations

AD: Alzheimer’s disease
APOE: Apolipoprotein E
cPLA_2_: cytosolic calcium-dependent phospholipase A_2_
AA: arachidonic acid
DHA: docosahexaenoic acid
EPA: eicosapentaenoic acid
iPLA_2_: calcium-independent phospholipase A_2_

