## Supplemental Information for "Development of Potent, Selective cPLA2 Inhibitors for Targeting Neuroinflammation in Alzheimer’s Disease and Other Neurodegenerative Disorders"

**Supplementary Table S1**. 1^St^ Generation compounds (117) selected by V-SYNTHES and synthesized for biochemical testing.

| **BRI-ID** | **Structure** | **Score** | **MW** | **Purity** | **Tanimoto distance to closest ChEMBL ligand** | **Closest ligand**  **ChEMBL ID** |
| --- | --- | --- | --- | --- | --- | --- |
| BRI-50000 | 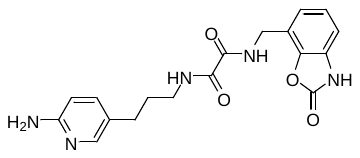 | -41.3 | 369.4 | 96 | 0.54 | CHEMBL3326956 |
| BRI-50001 | 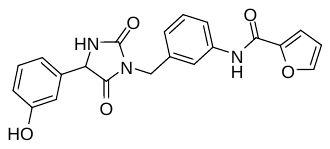 | -41.1 | 391.4 | 93 | 0.54 | CHEMBL428996 |
| BRI-50002 | 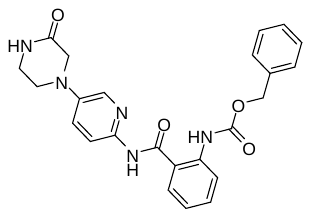 | -40.9 | 445.5 | 97 | 0.52 | CHEMBL9277 |
| BRI-50003 | 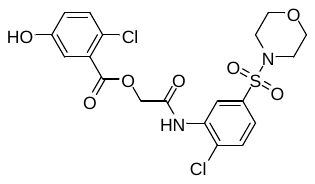 | -40.7 | 489.3 | 100 | 0.58 | CHEMBL267697 |
| BRI-50004 | 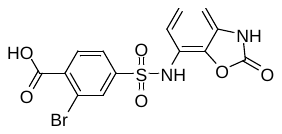 | -40.1 | 413.2 | 98 | 0.62 | CHEMBL408261 |
| BRI-50005 | 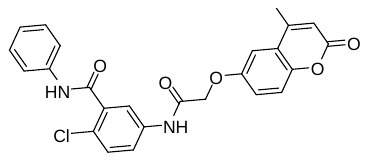 | -39.1 | 462.9 | 100 | 0.59 | CHEMBL199714 |
| BRI-50006 | 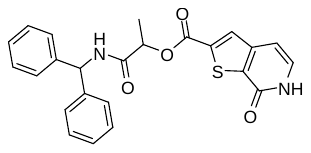 | -38.7 | 432.5 | 95 | 0.57 | CHEMBL268905 |
| BRI-50008 | 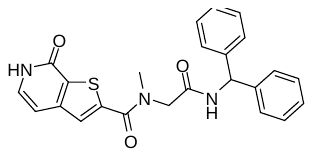 | -38.6 | 431.5 | 95 | 0.57 | CHEMBL268905 |
| BRI-50012 | 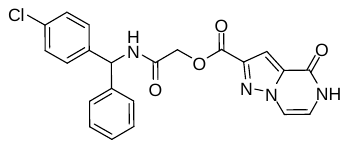 | -38.1 | 436.8 | 100 | 0.59 | CHEMBL375430 |
| BRI-50013 | 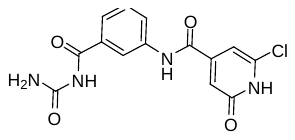 | -38.1 | 334.7 | 100 | 0.63 | CHEMBL8970 |
| BRI-50014 | 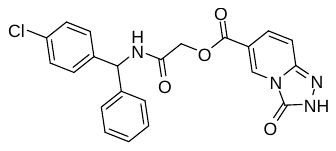 | -38.1 | 436.8 | 100 | 0.57 | CHEMBL375430 |
| BRI-50015 | 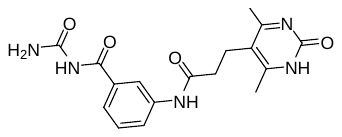 | -38.1 | 357.4 | 97 | 0.54 | CHEMBL112566 |
| BRI-50017 | 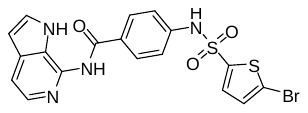 | -37.6 | 591.4 | 98 | 0.58 | CHEMBL345056 |
| BRI-50018 | 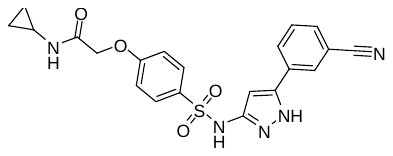 | -37.5 | 437.5 | 97 | 0.59 | CHEMBL242320 |
| BRI-50019 | 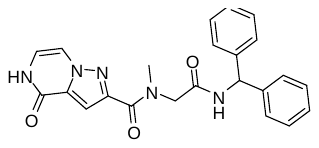 | -37.2 | 415.4 | 100 | 0.59 | CHEMBL268905 |
| BRI-50020 | 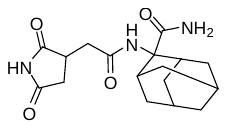 | -37.0 | 333.4 | 100 | 0.52 | CHEMBL95190 |
| BRI-50021 | 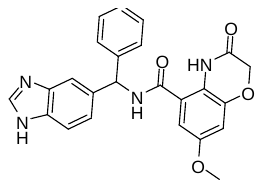 | -36.9 | 428.4 | 100 | 0.52 | CHEMBL242960 |
| BRI-50022 | 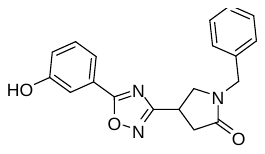 | -36.9 | 335.4 | 100 | 0.57 | CHEMBL208972 |
| BRI-50023 | 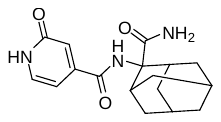 | -36.8 | 315.4 | 100 | 0.61 | CHEMBL428996 |
| BRI-50024 | 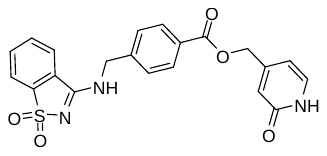 | -36.6 | 423.4 | 98 | 0.62 | CHEMBL428653 |
| BRI-50025 | 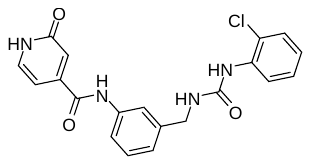 | -36.6 | 396.8 | 98 | 0.60 | CHEMBL8970 |
| BRI-50026 | 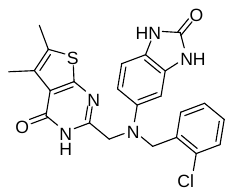 | -36.5 | 466.0 | 92 | 0.62 | CHEMBL375994 |
| BRI-50028 | 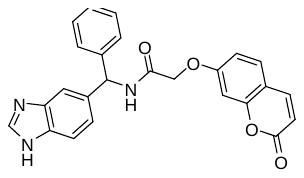 | -36.3 | 425.4 | 100 | 0.54 | CHEMBL242960 |
| BRI-50029 | 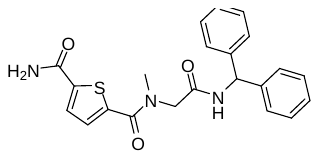 | -36.3 | 407.5 | 97 | 0.55 | CHEMBL428996 |
| BRI-50030 | 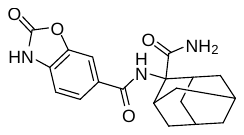 | -36.2 | 355.4 | 100 | 0.60 | CHEMBL242960 |
| BRI-50031 | 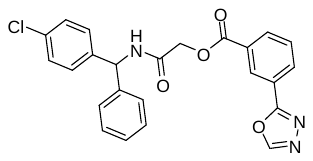 | -36.2 | 447.9 | 95 | 0.57 | CHEMBL210464 |
| BRI-50032 | 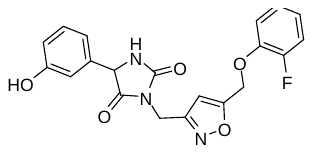 | -36.1 | 397.4 | 92 | 0.55 | CHEMBL268896 |
| BRI-50033 | 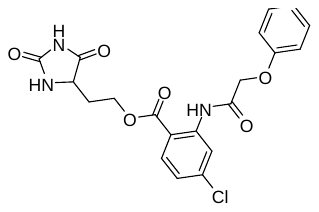 | -36.1 | 431.8 | 100 | 0.52 | CHEMBL416517 |
| BRI-50034 | 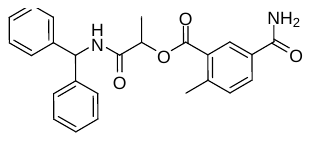 | -36.0 | 416.5 | 100 | 0.50 | CHEMBL268905 |
| BRI-50035 | 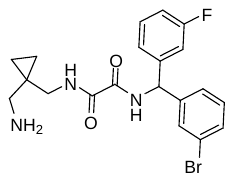 | -35.7 | 434.3 | 95 | 0.55 | CHEMBL428996 |
| BRI-50036 | 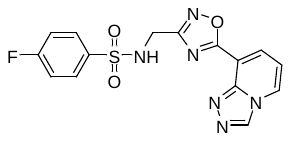 | -35.7 | 374.3 | 98 | 0.62 | CHEMBL408261 |
| BRI-50039 | 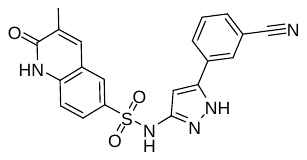 | -35.6 | 405.4 | 91 | 0.59 | CHEMBL242320 |
| BRI-50040 | 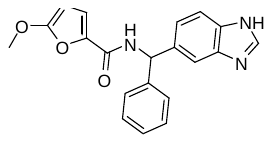 | -35.4 | 347.4 | 100 | 0.55 | CHEMBL242960 |
| BRI-50041 | 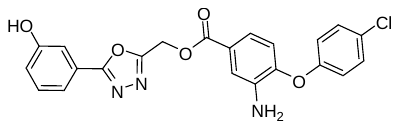 | -35.3 | 437.8 | 91 | 0.59 | CHEMBL609913 |
| BRI-50042 | 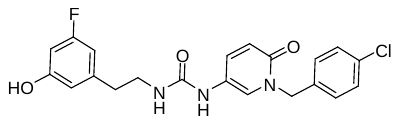 | -35.3 | 415.8 | 100 | 0.55 | CHEMBL375769 |
| BRI-50043 | 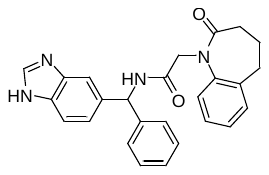 | -35.2 | 424.5 | 100 | 0.51 | CHEMBL242960 |
| BRI-50044 | 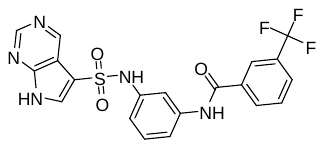 | -35.1 | 461.4 | 95 | 0.60 | CHEMBL345056 |
| BRI-50046 | 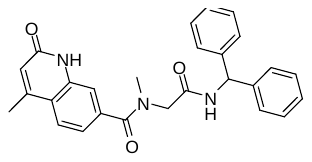 | -34.9 | 439.5 | 100 | 0.48 | CHEMBL242960 |
| BRI-50047 | 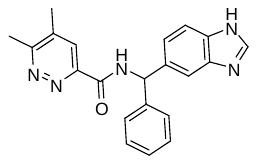 | -34.9 | 357.4 | 100 | 0.56 | CHEMBL337790 |
| BRI-50048 | 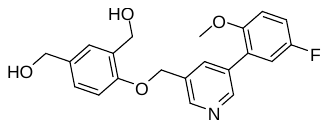 | -34.8 | 369.4 | 93 | 0.52 | CHEMBL598976 |
| BRI-50049 | 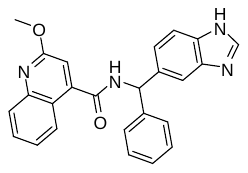 | -34.7 | 408.5 | 100 | 0.41 | CHEMBL344686 |
| BRI-50050 | 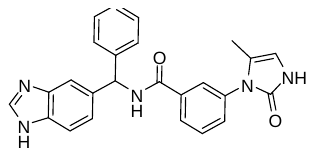 | -34.7 | 423.5 | 100 | 0.54 | CHEMBL3326949 |
| BRI-50051 | 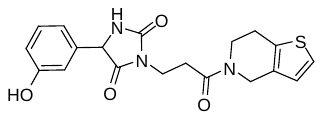 | -34.7 | 385.4 | 90 | 0.58 | CHEMBL428996 |
| BRI-50053 | 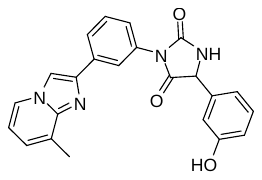 | -34.6 | 512.4 | 100 | 0.60 | CHEMBL3326967 |
| BRI-50054 | 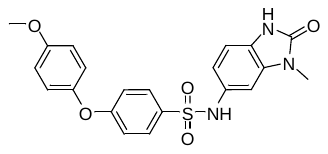 | -34.5 | 425.5 | 100 | 0.56 | CHEMBL220806 |
| BRI-50056 | 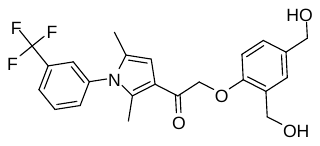 | -34.4 | 433.4 | 91 | 0.41 | CHEMBL598976 |
| BRI-50057 | 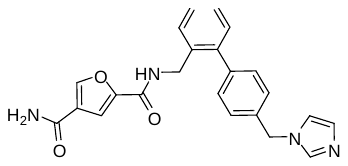 | -34.4 | 400.4 | 98 | 0.57 | CHEMBL210464 |
| BRI-50058 | 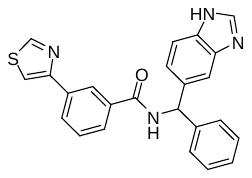 | -34.2 | 410.5 | 98 | 0.57 | CHEMBL138705 |
| BRI-50059 | 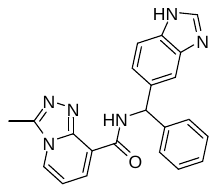 | -34.2 | 382.4 | 100 | 0.60 | CHEMBL201266 |
| BRI-50060 | 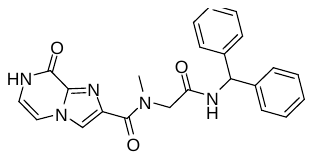 | -34.2 | 415.4 | 100 | 0.60 | CHEMBL428996 |
| BRI-50061 | 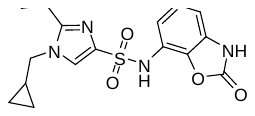 | -34.2 | 362.4 | 100 | 0.58 | CHEMBL463857 |
| BRI-50063 | 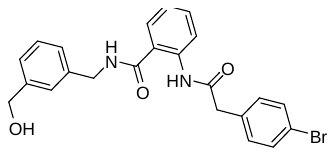 | -34.1 | 453.3 | 96 | 0.61 | CHEMBL164883 |
| BRI-50064 | 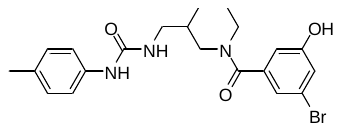 | -34.1 | 446.3 | 96 | 0.56 | CHEMBL428996 |
| BRI-50065 | 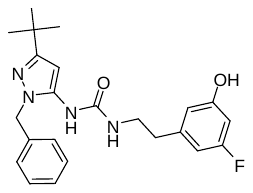 | -34.1 | 410.5 | 98 | 0.55 | CHEMBL609913 |
| BRI-50066 | 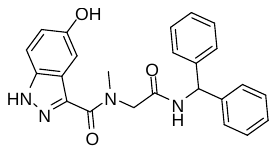 | -34.0 | 414.5 | 99 | 0.52 | CHEMBL242960 |
| BRI-50067 | 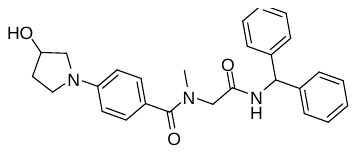 | -34.0 | 443.5 | 100 | 0.46 | CHEMBL268905 |
| BRI-50068 | 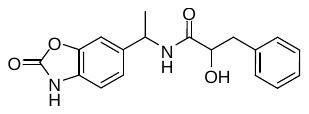 | -33.9 | 326.3 | 99 | 0.57 | CHEMBL242960 |
| BRI-50069 | 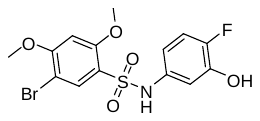 | -33.8 | 406.2 | 100 | 0.62 | CHEMBL408261 |
| BRI-50070 | 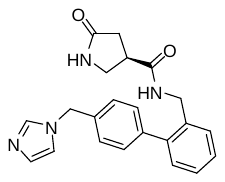 | -33.8 | 374.4 | 100 | 0.53 | CHEMBL598974 |
| BRI-50071 | 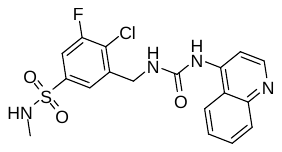 | -33.7 | 422.9 | 97 | 0.53 | CHEMBL408261 |
| BRI-50072 | 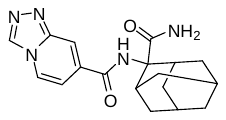 | -33.7 | 339.4 | 100 | 0.62 | CHEMBL242960 |
| BRI-50073 | 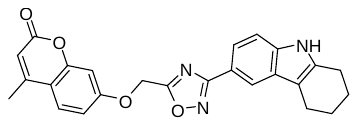 | -33.7 | 427.5 | 96 | 0.53 | CHEMBL241891 |
| BRI-50074 | 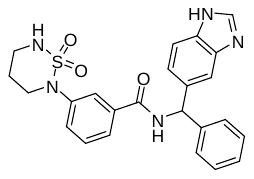 | -33.7 | 461.5 | 93 | 0.51 | CHEMBL375994 |
| BRI-50075 | 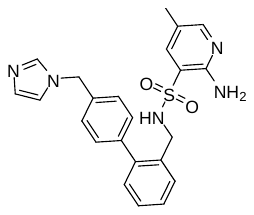 | -33.6 | 433.5 | 100 | 0.55 | CHEMBL463857 |
| BRI-50076 | 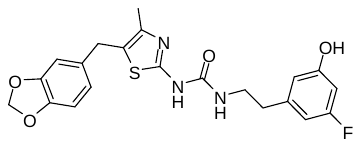 | -33.6 | 429.5 | 90 | 0.57 | CHEMBL375551 |
| BRI-50077 | 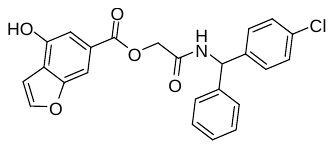 | -33.6 | 435.9 | 94 | 0.55 | CHEMBL375430 |
| BRI-50079 | 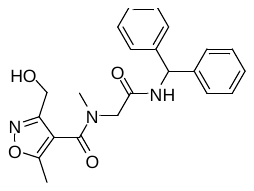 | -33.5 | 393.4 | 96 | 0.56 | CHEMBL268905 |
| BRI-50080 | 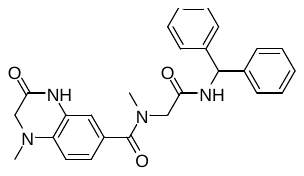 | -33.5 | 442.5 | 91 | 0.53 | CHEMBL242960 |
| BRI-50081 | 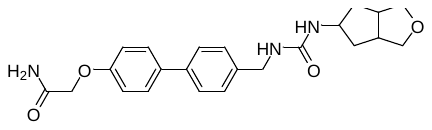 | -33.5 | 409.5 | 100 | 0.49 | CHEMBL30108 |
| BRI-50082 | 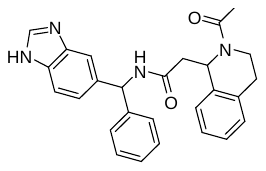 | -33.4 | 438.5 | 94 | 0.50 | CHEMBL3326956 |
| BRI-50083 | 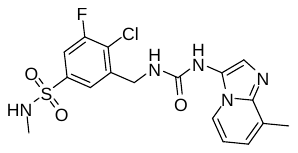 | -33.4 | 425.9 | 100 | 0.63 | CHEMBL370113 |
| BRI-50084 | 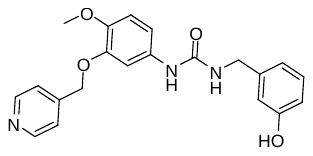 | -33.4 | 379.4 | 98 | 0.53 | CHEMBL598976 |
| BRI-50085 | 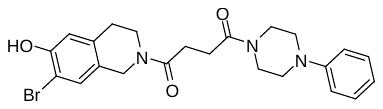 | -33.3 | 472.4 | 99 | 0.57 | CHEMBL428391 |
| BRI-50086 | 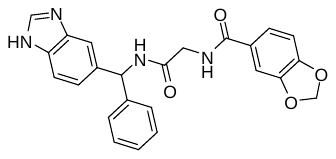 | -33.2 | 428.4 | 96 | 0.47 | CHEMBL242960 |
| BRI-50087 | 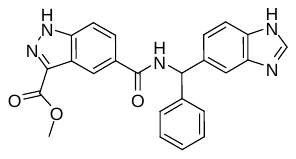 | -33.2 | 425.4 | 100 | 0.57 | CHEMBL344686 |
| BRI-50089 | 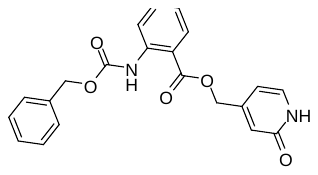 | -33.1 | 378.4 | 100 | 0.57 | CHEMBL268905 |
| BRI-50090 | 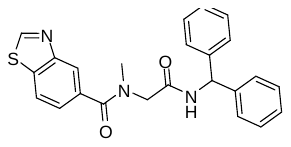 | -33.1 | 415.5 | 98 | 0.52 | CHEMBL242960 |
| BRI-50091 | 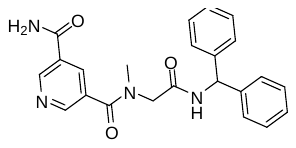 | -33.1 | 402.4 | 100 | 0.54 | CHEMBL428996 |
| BRI-50092 | 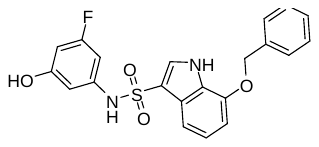 | -33.0 | 412.4 | 100 | 0.53 | CHEMBL220806 |
| BRI-50093 | 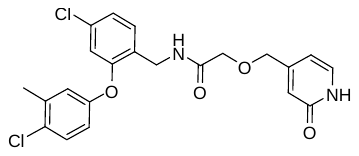 | -33.0 | 447.3 | 100 | 0.55 | CHEMBL428827 |
| BRI-50095 | 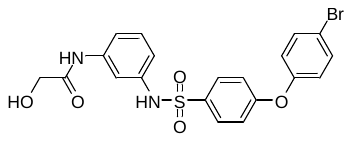 | -32.9 | 477.3 | 90 | 0.60 | CHEMBL345056 |
| BRI-50096 | 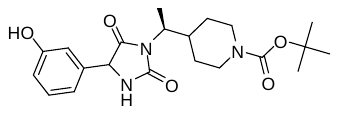 | -32.9 | 403.5 | 99 | 0.52 | CHEMBL428996 |
| BRI-50097 | 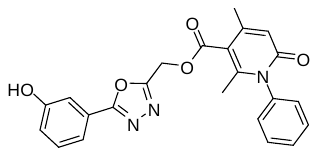 | -32.8 | 417.4 | 100 | 0.57 | CHEMBL3326971 |
| BRI-50098 | 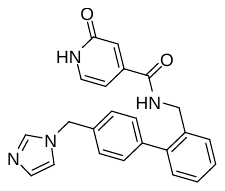 | -32.8 | 384.4 | 100 | 0.54 | CHEMBL208073 |
| BRI-50099 | 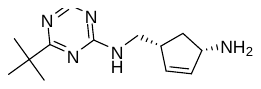 | -32.8 | 247.3 | 91 | 0.71 | CHEMBL207086 |
| BRI-50100 | 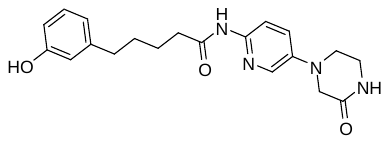 | -32.6 | 368.4 | 100 | 0.58 | CHEMBL3326956 |
| BRI-50101 | 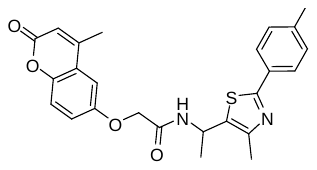 | -32.6 | 448.5 | 98 | 0.55 | CHEMBL3329869 |
| BRI-50102 | 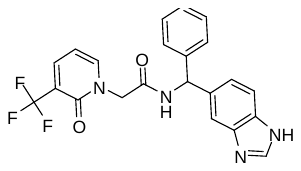 | -32.6 | 426.4 | 100 | 0.53 | CHEMBL3326965 |
| BRI-50103 | 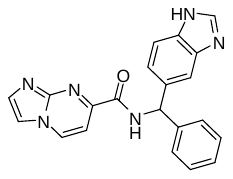 | -32.6 | 368.4 | 97 | 0.61 | CHEMBL242960 |
| BRI-50106 | 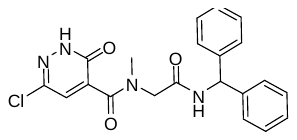 | -32.5 | 410.9 | 97 | 0.58 | CHEMBL268905 |
| BRI-50107 | 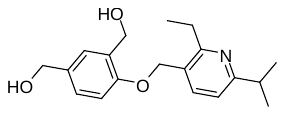 | -32.5 | 315.4 | 100 | 0.47 | CHEMBL598976 |
| BRI-50108 | 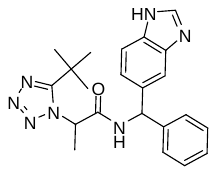 | -32.4 | 403.5 | 100 | 0.55 | CHEMBL509739 |
| BRI-50109 | 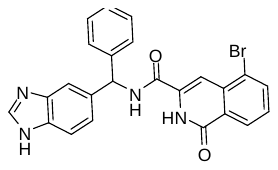 | -32.4 | 473.3 | 100 | 0.55 | CHEMBL375430 |
| BRI-50110 | 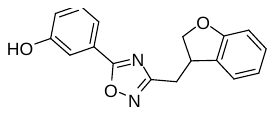 | -32.4 | 294.3 | 91 | 0.54 | CHEMBL241891 |
| BRI-50111 | 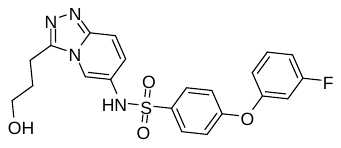 | -32.4 | 442.5 | 98 | 0.60 | CHEMBL220806 |
| BRI-50112 | 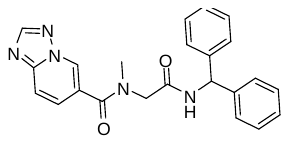 | -32.3 | 399.4 | 100 | 0.60 | CHEMBL242960 |
| BRI-50113 | 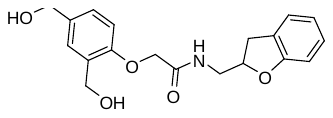 | -32.3 | 343.4 | 91 | 0.52 | CHEMBL428391 |
| BRI-50114 | 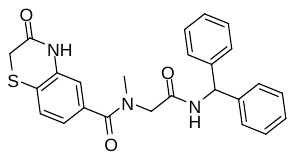 | -32.2 | 445.5 | 100 | 0.53 | CHEMBL242960 |
| BRI-50115 | 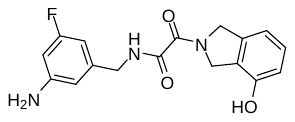 | -32.2 | 329.3 | 93 | 0.61 | CHEMBL428996 |
| BRI-50116 | 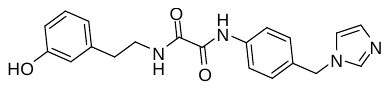 | -32.2 | 364.4 | 100 | 0.48 | CHEMBL598974 |
| BRI-50117 |  | -32.1 | 483.3 | 97 | 0.58 | CHEMBL8970 |
| BRI-50118 |  | -32.0 | 423.5 | 100 | 0.62 | CHEMBL463857 |
| BRI-50119 |  | -32.0 | 346.4 | 91 | 0.54 | CHEMBL408261 |
| BRI-50120 |  | -31.9 | 373.4 | 100 | 0.54 | CHEMBL345056 |
| BRI-50121 |  | -31.9 | 410.3 | 100 | 0.63 | CHEMBL139508 |
| BRI-50122 |  | -31.8 | 410.4 | 100 | 0.58 | CHEMBL428996 |
| BRI-50123 |  | -31.7 | 399.4 | 100 | 0.52 | CHEMBL138705 |
| BRI-50124 |  | -31.6 | 427.5 | 96 | 0.46 | CHEMBL609913 |
| BRI-50125 |  | -31.6 | 417.3 | 94 | 0.58 | CHEMBL3326961 |
| BRI-50126 |  | -31.6 | 473.3 | 98 | 0.54 | CHEMBL408261 |
| BRI-50127 |  | -31.3 | 474.0 | 97 | 0.44 | CHEMBL428391 |
| BRI-50128 |  | -31.1 | 340.4 | 100 | 0.60 | CHEMBL508499 |
| BRI-50129 |  | -31.0 | 465.3 | 100 | 0.55 | CHEMBL375769 |
| BRI-50130 |  | -30.8 | 388.5 | 100 | 0.50 | CHEMBL201266 |
| BRI-50131 |  | -30.7 | 327.4 | 92 | 0.64 | CHEMBL207418 |
| BRI-50132 |  | -30.6 | 379.3 | 100 | 0.55 | CHEMBL344686 |
| BRI-50134 |  | -29.9 | 401.5 | 100 | 0.58 | CHEMBL504084 |

**Supplementary Table S2.** Dose-response curves for hit compounds from the *in vitro* cPLA_2_ fluorescence assay.

| **Cmpd.** | **Structure** | **Dose-response data** | **IC_50_ (µM)** |
| --- | --- | --- | --- |
| ASB-14780 |  |  | 6.66 |
| BRI-50006 |  |  | 5.35 |
| BRI-50026 |  |  | 8.30 |
| BRI-50031 |  |  | 8.3 |
| BRI-50049 |  |  | 2.32 |
| BRI-50054 |  |  | 3.22 |
| BRI-50077 |  |  | 4.43 |
| BRI-50125 |  |  | 3.18 |

| nM inhibitor | y |
| --- | --- |
| 9.77 | 112.412 |
| 9.77 | 112.463 |
| 39.1 | 103.846 |
| 39.1 | 113.045 |
| 625 | 102.035 |
| 625 | 98.3403 |
| 2500 | 85.5279 |
| 2500 | 85.5636 |
| 10000 | 31.6347 |
| 10000 | 32.7938 |

| nM inhibitor | % Relative Activity |
| --- | --- |
| 1 | 100 |
| 2 | 100 |
| 4 | 99.9999 |
| 8 | 99.9996 |
| 16 | 99.9985 |
| 32 | 99.9946 |
| 64 | 99.9807 |
| 128 | 99.9309 |
| 256 | 99.7529 |
| 512 | 99.1203 |
| 1024 | 96.9175 |
| 2048 | 89.7683 |
| 4096 | 70.9996 |
| 8192 | 40.5884 |
| 16384 | 16.0114 |
| 32768 | 5.05101 |
| 65536 | 1.46274 |
| 131072 | 0.412524 |

| nM inhibitor | % Relative Activity |
| --- | --- |
| 1 | 99.9989 |
| 2 | 99.9972 |
| 4 | 99.9931 |
| 8 | 99.9826 |
| 16 | 99.9563 |
| 32 | 99.89 |
| 64 | 99.7239 |
| 128 | 99.3086 |
| 256 | 98.2791 |
| 512 | 95.782 |
| 1024 | 90.0289 |
| 2048 | 78.2139 |
| 4096 | 58.8049 |
| 8192 | 36.2077 |
| 16384 | 18.4128 |
| 32768 | 8.23457 |
| 65536 | 3.44509 |
| 131072 | 1.39885 |

| nM inhibitor | y |
| --- | --- |
| 9.77 | 95.7458 |
| 9.77 | 93.944 |
| 39.1 | 106.031 |
| 39.1 | 103.872 |
| 625 | 100.042 |
| 625 | 95.2376 |
| 2500 | 73.8305 |
| 2500 | 69.4186 |
| 10000 | 32.9991 |
| 10000 | 29.313 |

| nM inhibitor | y |
| --- | --- |
| 9.77 | 122.022 |
| 9.77 | 120.823 |
| 39.1 | 127.493 |
| 39.1 | 123.752 |
| 625 | 125.376 |
| 625 | 117.757 |
| 2500 | 99.1838 |
| 2500 | 97.0696 |
| 10000 | 34.9555 |
| 10000 | 33.711 |

| nM inhibitor | % Relative Activity |
| --- | --- |
| 1 | 100 |
| 2 | 100 |
| 4 | 100 |
| 8 | 100 |
| 16 | 100 |
| 32 | 100 |
| 64 | 100 |
| 128 | 100 |
| 256 | 99.9994 |
| 512 | 99.9938 |
| 1024 | 99.9311 |
| 2048 | 99.2369 |
| 4096 | 92.1026 |
| 8192 | 51.1205 |
| 16384 | 8.57461 |
| 32768 | 0.834045 |
| 65536 | 0.0753666 |
| 131072 | 0.00676327 |

| nM inhibitor | y |
| --- | --- |
| 9.77 | 107.361 |
| 9.77 | 103.289 |
| 39.1 | 93.2021 |
| 39.1 | 88.0497 |
| 625 | 104.906 |
| 625 | 97.8553 |
| 2500 | 98.8594 |
| 2500 | 91.9825 |
| 10000 | 48.0512 |
| 10000 | 44.0281 |

| nM inhibitor | % Relative Activity |
| --- | --- |
| 1 | 100 |
| 2 | 100 |
| 4 | 100 |
| 8 | 100 |
| 16 | 100 |
| 32 | 100 |
| 64 | 100 |
| 128 | 100 |
| 256 | 99.9994 |
| 512 | 99.9938 |
| 1024 | 99.9311 |
| 2048 | 99.2369 |
| 4096 | 92.1026 |
| 8192 | 51.1204 |
| 16384 | 8.57457 |
| 32768 | 0.83404 |
| 65536 | 0.075366 |
| 131072 | 0.0067632 |

| nM inhibitor | y |
| --- | --- |
| 9.77 | 99.7569 |
| 9.77 | 99.378 |
| 39.1 | 96.2682 |
| 39.1 | 94.3947 |
| 625 | 81.6413 |
| 625 | 79.1589 |
| 2500 | 40.1977 |
| 2500 | 38.6584 |
| 10000 | 30.8211 |
| 10000 | 28.4856 |

| nM inhibitor | % Relative Activity |
| --- | --- |
| 1 | 99.8079 |
| 2 | 99.6643 |
| 4 | 99.4142 |
| 8 | 98.9796 |
| 16 | 98.2282 |
| 32 | 96.9408 |
| 64 | 94.7677 |
| 128 | 91.1913 |
| 256 | 85.5431 |
| 512 | 77.1797 |
| 1024 | 65.9062 |
| 2048 | 52.4918 |
| 4096 | 38.7078 |
| 8192 | 26.5227 |
| 16384 | 17.103 |
| 32768 | 10.5485 |
| 65536 | 6.31462 |
| 131072 | 3.70962 |

| nM inhibitor | y |
| --- | --- |
| 9.77 | 86.3339 |
| 9.77 | 84.5593 |
| 156 | 89.5285 |
| 156 | 88.808 |
| 625 | 79.2855 |
| 625 | 74.213 |
| 2500 | 60.6922 |
| 2500 | 58.5416 |
| 10000 | 27.1245 |
| 10000 | 24.7355 |

| nM inhibitor | % Relative Activity |
| --- | --- |
| 1 | 99.7325 |
| 2 | 99.5562 |
| 4 | 99.2646 |
| 8 | 98.7835 |
| 16 | 97.9942 |
| 32 | 96.7098 |
| 64 | 94.6478 |
| 128 | 91.4084 |
| 256 | 86.4882 |
| 512 | 79.3857 |
| 1024 | 69.8513 |
| 2048 | 58.2276 |
| 4096 | 45.6117 |
| 8192 | 33.5349 |
| 16384 | 23.2867 |
| 32768 | 15.4426 |
| 65536 | 9.89981 |
| 131072 | 6.2006 |

| nM inhibitor | y |
| --- | --- |
| 9.77 | 84.222 |
| 9.77 | 84.7597 |
| 156 | 95.3299 |
| 156 | 90.4928 |
| 625 | 83.9176 |
| 625 | 80.8901 |
| 2500 | 66.2204 |
| 2500 | 62.4567 |
| 10000 | 32.7138 |
| 10000 | 30.9613 |

| nM inhibitor | % Relative Activity |
| --- | --- |
| 1 | 99.8772 |
| 2 | 99.7867 |
| 4 | 99.6296 |
| 8 | 99.3577 |
| 16 | 98.8884 |
| 32 | 98.0829 |
| 64 | 96.713 |
| 128 | 94.4198 |
| 256 | 90.6811 |
| 512 | 84.8395 |
| 1024 | 76.2935 |
| 2048 | 64.9219 |
| 4096 | 51.5589 |
| 8192 | 37.9692 |
| 16384 | 26.0362 |
| 32768 | 16.8357 |
| 65536 | 10.428 |
| 131072 | 6.27506 |

| nM inhibitor | y |
| --- | --- |
| 9.77 | 91.4522 |
| 9.77 | 91.0681 |
| 156 | 92.2947 |
| 156 | 89.9663 |
| 625 | 91.7977 |
| 625 | 88.9874 |
| 2500 | 53.2422 |
| 2500 | 50.0196 |
| 10000 | 25.4092 |
| 10000 | 27.0127 |

| nM inhibitor | %Relative Activity |
| --- | --- |
| 1 | 99.9712 |
| 2 | 99.942 |
| 4 | 99.8833 |
| 8 | 99.7651 |
| 16 | 99.5278 |
| 32 | 99.0529 |
| 64 | 98.1096 |
| 128 | 96.2623 |
| 256 | 92.7433 |
| 512 | 86.3801 |
| 1024 | 75.8877 |
| 2048 | 60.9649 |
| 4096 | 43.6629 |
| 8192 | 27.7771 |
| 16384 | 16.0268 |
| 32768 | 8.65165 |
| 65536 | 4.48896 |
| 131072 | 2.27915 |

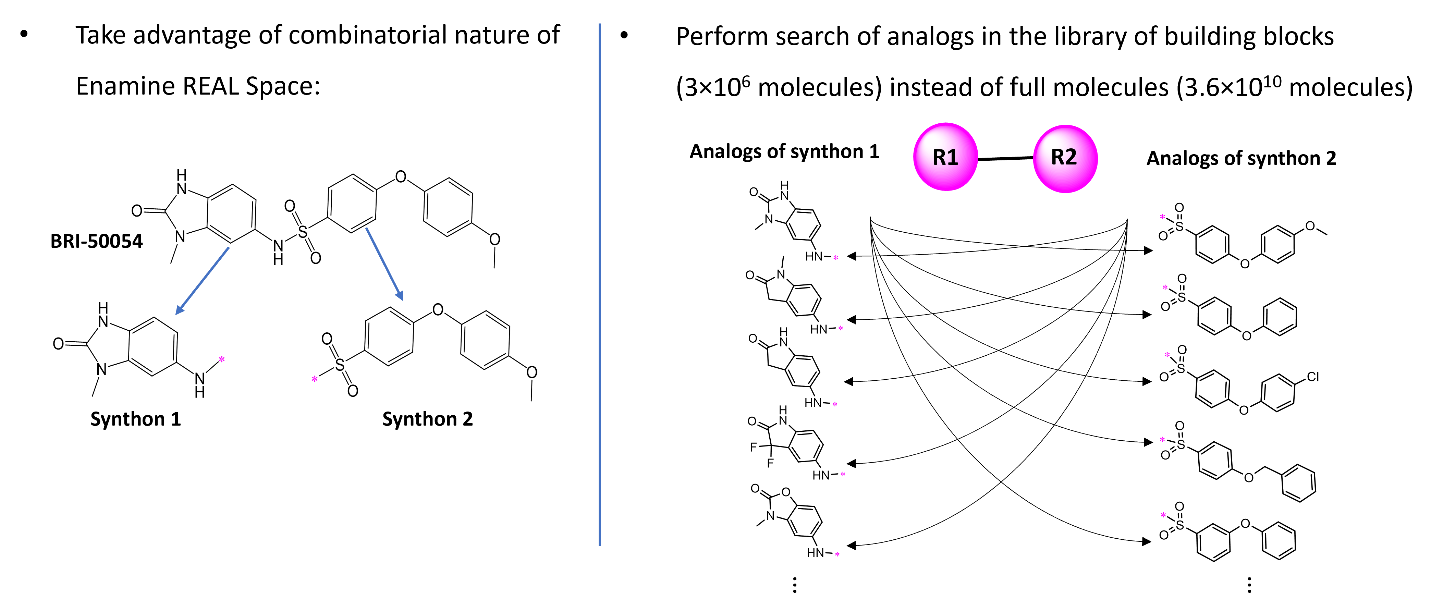
**Supplementary Figure S1:** An example of search algorithm performing decomposition of hit molecule and search for analogs in the whole REAL Space without enumeration of all molecules.

**Supplementary Table S3.** Analogs of compounds BRI-50054, BRI-50077 and BRI-50125 selected for testing on the first round of optimization.

| **BRI-ID** | **Structure** | **Score** | **MW** | **Purity** | **Tanimoto distance to**  **closest ChEMBL ligand** | **Closest ligand ChEMBL ID** |
| --- | --- | --- | --- | --- | --- | --- |
| BRI-50054 analogs | | | | | | |
| BRI-50200 |  | -34.6 | 425.5 | 90 | 0.57 | CHEMBL242320 |
| BRI-50201 |  | -34.6 | 474.3 | 99 | 0.58 | CHEMBL242320 |
| BRI-50202 |  | -34.2 | 410.4 | 100 | 0.50 | CHEMBL242320 |
| BRI-50203 |  | -34.1 | 446.9 | 98 | 0.57 | CHEMBL3327096 |
| BRI-50204 |  | -33.5 | 456.5 | 100 | 0.59 | CHEMBL345056 |
| BRI-50206 |  | -33.2 | 422.5 | 100 | 0.59 | CHEMBL345056 |
| BRI-50207 |  | -32.7 | 436.5 | 98 | 0.52 | CHEMBL220806 |
| BRI-50208 |  | -32.7 | 459.3 | 95 | 0.51 | CHEMBL242320 |
| BRI-50209 |  | -32.7 | 482.3 | 93 | 0.57 | CHEMBL220806 |
| BRI-50210 |  | -32.5 | 450.5 | 100 | 0.46 | CHEMBL220806 |
| BRI-50212 |  | -32.2 | 438.5 | 96 | 0.51 | CHEMBL345056 |
| BRI-50214 |  | -32.2 | 455.9 | 100 | 0.48 | CHEMBL520836 |
| BRI-50215 |  | -32.0 | 436.5 | 96 | 0.55 | CHEMBL345056 |
| BRI-50217 |  | -32.0 | 457.3 | 94 | 0.55 | CHEMBL408261 |
| BRI-50218 |  | -31.8 | 429.9 | 100 | 0.56 | CHEMBL220806 |
| BRI-50219 |  | -31.6 | 437.9 | 100 | 0.56 | CHEMBL220806 |
| BRI-50220 |  | -31.6 | 471.4 | 100 | 0.55 | CHEMBL408261 |
| BRI-50221 |  | -31.6 | 424.5 | 98 | 0.50 | CHEMBL345056 |
| BRI-50224 |  | -31.3 | 422.5 | 100 | 0.62 | CHEMBL210464 |
| BRI-50226 |  | -31.0 | 423.4 | 92 | 0.60 | CHEMBL345056 |
| BRI-50227 |  | -30.8 | 518.4 | 100 | 0.49 | CHEMBL520836 |
| BRI-50228 |  | -30.6 | 452.5 | 100 | 0.50 | CHEMBL345056 |
| BRI-50229 |  | -30.3 | 442.5 | 100 | 0.56 | CHEMBL345056 |
| BRI-50230 |  | -30.3 | 476.9 | 98 | 0.56 | CHEMBL408261 |
| BRI-50232 |  | -27.8 | 430.9 | 92 | 0.57 | CHEMBL220806 |
| BRI-50233 |  | -26.7 | 430.9 | 100 | 0.57 | CHEMBL220806 |
| BRI-50236 |  | -23.1 | 430.5 | 100 | 0.58 | CHEMBL345056 |
| BRI-50237 |  | -23.0 | 439.5 | 98 | 0.55 | CHEMBL520836 |
| BRI-50238 |  | -22.8 | 440.5 | 100 | 0.57 | CHEMBL242320 |
| BRI-50077 analogs | | | | | | |
| BRI-50239 |  | -34.3 | 440.9 | 100 | 0.52 | CHEMBL8970 |
| BRI-50240 |  | -34.2 | 415.4 | 98 | 0.57 | CHEMBL242960 |
| BRI-50241 |  | -34.2 | 422.9 | 98 | 0.55 | CHEMBL428996 |
| BRI-50243 |  | -33.3 | 461.5 | 91 | 0.53 | CHEMBL242960 |
| BRI-50244 |  | -33.1 | 388.5 | 100 | 0.53 | CHEMBL428996 |
| BRI-50245 |  | -32.8 | 429.3 | 100 | 0.55 | CHEMBL29838 |
| BRI-50246 |  | -32.4 | 439.3 | 99 | 0.55 | CHEMBL29838 |
| BRI-50247 |  | -31.8 | 435.9 | 100 | 0.55 | CHEMBL375430 |
| BRI-50248 |  | -31.8 | 401.4 | 100 | 0.56 | CHEMBL375430 |
| BRI-50249 |  | -31.7 | 477.9 | 95 | 0.54 | CHEMBL375430 |
| BRI-50251 |  | -31.6 | 435.5 | 100 | 0.57 | CHEMBL242960 |
| BRI-50252 |  | -31.6 | 437.5 | 100 | 0.57 | CHEMBL242960 |
| BRI-50254 |  | -31.5 | 457.5 | 95 | 0.55 | CHEMBL242960 |
| BRI-50255 |  | -31.4 | 414.5 | 100 | 0.56 | CHEMBL242960 |
| BRI-50256 |  | -31.4 | 407.4 | 91 | 0.58 | CHEMBL242960 |
| BRI-50257 |  | -31.1 | 373.8 | 100 | 0.56 | CHEMBL375430 |
| BRI-50258 |  | -30.9 | 373.8 | 100 | 0.56 | CHEMBL375430 |
| BRI-50259 |  | -30.8 | 431.4 | 91 | 0.53 | CHEMBL242960 |
| BRI-50260 |  | -30.7 | 392.4 | 99 | 0.51 | CHEMBL428996 |
| BRI-50261 |  | -30.5 | 415.4 | 97 | 0.55 | CHEMBL242960 |
| BRI-50262 |  | -30.2 | 387.8 | 100 | 0.55 | CHEMBL242960 |
| BRI-50263 |  | -30.0 | 443.4 | 100 | 0.54 | CHEMBL138867 |
| BRI-50264 |  | -29.9 | 408.4 | 100 | 0.55 | CHEMBL3329874 |
| BRI-50265 |  | -29.8 | 415.5 | 100 | 0.52 | CHEMBL242960 |
| BRI-50266 |  | -29.6 | 360.4 | 100 | 0.61 | CHEMBL242960 |
| BRI-50267 |  | -29.6 | 398.5 | 100 | 0.58 | CHEMBL242960 |
| BRI-50268 |  | -29.1 | 361.3 | 100 | 0.62 | CHEMBL443834 |
| BRI-50125 analogs | | | | | | |
| BRI-50269 |  | -36.9 | 507.5 | 100 | 0.54 | CHEMBL242960 |
| BRI-50271 |  | -35.4 | 415.4 | 100 | 0.53 | CHEMBL205219 |
| BRI-50272 |  | -35.3 | 439.5 | 99 | 0.52 | CHEMBL3326961 |
| BRI-50273 |  | -34.9 | 477.4 | 100 | 0.55 | CHEMBL199533 |
| BRI-50274 |  | -34.9 | 403.4 | 100 | 0.59 | CHEMBL3326961 |
| BRI-50276 |  | -33.8 | 496.3 | 100 | 0.58 | CHEMBL199533 |
| BRI-50277 |  | -33.8 | 436.8 | 100 | 0.58 | CHEMBL199714 |
| BRI-50278 |  | -33.7 | 391.4 | 100 | 0.58 | CHEMBL3326961 |
| BRI-50279 |  | -33.3 | 403.4 | 100 | 0.58 | CHEMBL3326961 |
| BRI-50280 |  | -33.2 | 451.9 | 95 | 0.57 | CHEMBL199533 |
| BRI-50281 |  | -32.8 | 457.5 | 98 | 0.56 | CHEMBL3326961 |
| BRI-50282 |  | -32.4 | 440.5 | 100 | 0.56 | CHEMBL3326961 |
| BRI-50283 |  | -32.4 | 495.3 | 100 | 0.59 | CHEMBL3326961 |
| BRI-50284 |  | -32.1 | 451.9 | 98 | 0.57 | CHEMBL199533 |
| BRI-50285 |  | -31.1 | 425.4 | 100 | 0.54 | CHEMBL3326961 |
| BRI-50286 |  | -31.0 | 430.5 | 96 | 0.56 | CHEMBL3326961 |
| BRI-50287 |  | -31.0 | 409.8 | 100 | 0.59 | CHEMBL3326961 |
| BRI-50288 |  | -31.0 | 391.4 | 91 | 0.59 | CHEMBL205219 |
| BRI-50289 |  | -30.6 | 465.9 | 100 | 0.57 | CHEMBL205219 |
| BRI-50290 |  | -30.6 | 402.4 | 100 | 0.55 | CHEMBL3326961 |
| BRI-50291 |  | -30.4 | 375.4 | 100 | 0.58 | CHEMBL3326961 |
| BRI-50292 |  | -30.0 | 375.4 | 100 | 0.56 | CHEMBL3326961 |
| BRI-50293 |  | -30.0 | 376.4 | 97 | 0.58 | CHEMBL205219 |
| BRI-50294 |  | -30.0 | 426.8 | 100 | 0.59 | CHEMBL205219 |
| BRI-50295 |  | -30.0 | 451.9 | 91 | 0.57 | CHEMBL199533 |
| BRI-50296 |  | -29.9 | 390.4 | 98 | 0.61 | CHEMBL3326961 |
| BRI-50297 |  | -26.3 | 362.4 | 95 | 0.50 | CHEMBL3326961 |
| BRI-50298 |  | -25.9 | 362.4 | 100 | 0.49 | CHEMBL3326961 |
| BRI-50299 |  | -25.1 | 376.4 | 100 | 0.49 | CHEMBL3326961 |

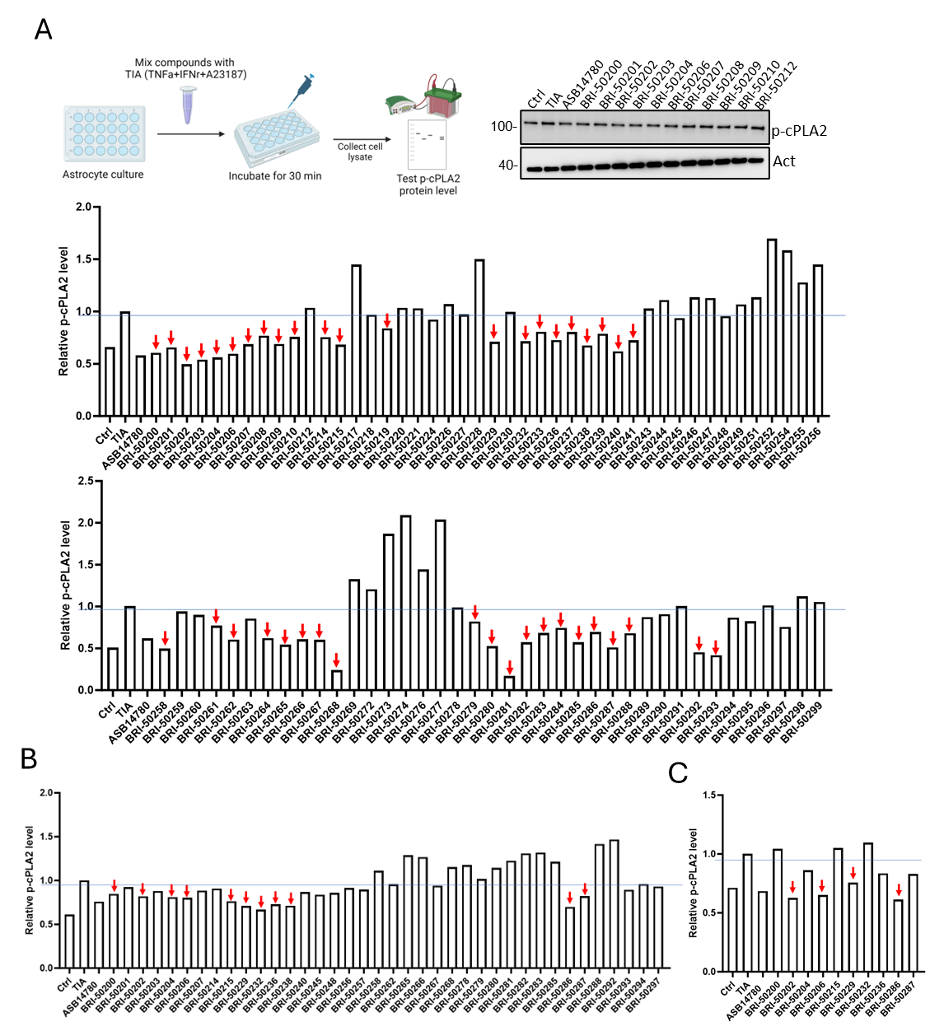

**Supplementary Figure S2:** Screening of 2^nd^ Generation compounds on the inhibitory effect of cPLA_2_ phosphorylation using in vitro assay. **(A)** Diagram of the assay and representative image of Western blot. The first round of screening consisted of eighty-three compounds (10 μM) that were tested for an inhibitory effect on cPLA_2_ phosphorylation. Compounds showing inhibitory effects on cPLA_2_ phosphorylation are indicated by arrows. **(B)** Thirty-nine compounds (5 μM) from the first round of screening were tested for their inhibitory effect on cPLA_2_ phosphorylation. Compounds showing inhibitory effects on cPLA_2_ phosphorylation are indicated again by arrows. **(C)** Inhibitory effects on cPLA_2_ phosphorylation were tested on ten compounds (1 μM) from the second round of screening. Compounds showing inhibitory effects on cPLA_2_ phosphorylation are indicated by an arrowhead.

**Supplementary Table S4.** Analogs of compounds BRI-50202 and BRI-50281 selected for testing on the second round of optimization.

| **BRI-ID** | **Structure** | **Score** | **MW** | **Purity** | **Tanimoto distance to**  **Closest ChEMBL ligand** | **Closest ligand ChEMBL ID** |
| --- | --- | --- | --- | --- | --- | --- |
| BRI-50202 analogs | | | | | | |
| BRI-50300 |  | -31.6 | 473.5 | 91 | 0.58 | CHEMBL220806 |
| BRI-50301 |  | -30.3 | 424.5 | 100 | 0.53 | CHEMBL408261 |
| BRI-50303 |  | -30.1 | 477.3 | 100 | 0.52 | CHEMBL242320 |
| BRI-50305 |  | -28.7 | 426.4 | 100 | 0.58 | CHEMBL345056 |
| BRI-50306 |  | -28.3 | 429.4 | 100 | 0.52 | CHEMBL29838 |
| BRI-50308 |  | -28.2 | 410.4 | 100 | 0.50 | CHEMBL345056 |
| BRI-50313 |  | -27.6 | 424.5 | 97 | 0.48 | CHEMBL242320 |
| BRI-50314 |  | -27.3 | 412.4 | 94 | 0.60 | CHEMBL345056 |
| BRI-50315 |  | -26.9 | 414.9 | 100 | 0.50 | CHEMBL220806 |
| BRI-50316 |  | -26.9 | 394.4 | 91 | 0.48 | CHEMBL220806 |
| BRI-50318 |  | -26.7 | 412.4 | 100 | 0.59 | CHEMBL345056 |
| BRI-50320 |  | -26.2 | 412.4 | 95 | 0.50 | CHEMBL242320 |
| BRI-50321 |  | -26.1 | 438.5 | 100 | 0.57 | CHEMBL345056 |
| BRI-50323 |  | -25.8 | 382.2 | 95 | 0.61 | CHEMBL220806 |
| BRI-50324 |  | -25.3 | 426.4 | 100 | 0.57 | CHEMBL345056 |
| BRI-50329 |  | -24.8 | 438.5 | 98 | 0.54 | CHEMBL220577 |
| BRI-50331 |  | -24.6 | 405.8 | 91 | 0.58 | CHEMBL220806 |
| BRI-50333 |  | -24.4 | 367.8 | 100 | 0.57 | CHEMBL220806 |
| BRI-50335 |  | -24.2 | 394.4 | 100 | 0.49 | CHEMBL345056 |
| BRI-50336 |  | -24.2 | 398.4 | 100 | 0.51 | CHEMBL220806 |
| BRI-50340 |  | -23.2 | 346.4 | 100 | 0.49 | CHEMBL242320 |
| BRI-50342 |  | -23.0 | 346.4 | 100 | 0.48 | CHEMBL345056 |
| BRI-50345 |  | -21.9 | 376.4 | 97 | 0.47 | CHEMBL242320 |
| BRI-50347 |  | -20.8 | 392.4 | 100 | 0.50 | CHEMBL345056 |
| BRI-50348 |  | -20.1 | 380.4 | 100 | 0.51 | CHEMBL220806 |
| BRI-50349 |  | -19.0 | 408.5 | 100 | 0.47 | CHEMBL345056 |
| BRI-50350 |  | -18.4 | 374.5 | 100 | 0.45 | CHEMBL242320 |
| BRI-50352 |  | -17.5 | 362.4 | 97 | 0.48 | CHEMBL242320 |
| BRI-50353 |  | -17.5 | 398.4 | 100 | 0.51 | CHEMBL242320 |
| BRI-50354 |  | -17.5 | 350.4 | 100 | 0.50 | CHEMBL242320 |
| BRI-50357 |  | -15.4 | 394.4 | 100 | 0.49 | CHEMBL345056 |
| BRI-50358 |  | -15.2 | 380.4 | 100 | 0.50 | CHEMBL345056 |
| BRI-50360 |  | -14.5 | 380.4 | 100 | 0.51 | CHEMBL242320 |
| BRI-50363 |  | -13.3 | 398.4 | 100 | 0.51 | CHEMBL242320 |
| BRI-50365 |  | -12.8 | 368.4 | 91 | 0.50 | CHEMBL242320 |
| BRI-50368 |  | -11.5 | 347.4 | 100 | 0.55 | CHEMBL220806 |
| BRI-50372 |  | -10.6 | 359.4 | 91 | 0.54 | CHEMBL345056 |
| BRI-50373 |  | -10.5 | 375.4 | 100 | 0.54 | CHEMBL220806 |
| BRI-50376 |  | -10.3 | 346.4 | 100 | 0.47 | CHEMBL242320 |
| BRI-50377 |  | -10.2 | 365.4 | 100 | 0.55 | CHEMBL220806 |
| BRI-50380 |  | -9.8 | 332.4 | 100 | 0.49 | CHEMBL242320 |
| BRI-50381 |  | -9.5 | 394.4 | 91 | 0.48 | CHEMBL242320 |
| BRI-50383 |  | -9.4 | 361.4 | 100 | 0.55 | CHEMBL220806 |
| BRI-50385 |  | -9.1 | 360.4 | 91 | 0.47 | CHEMBL242320 |
| BRI-50386 |  | -8.9 | 386.3 | 98 | 0.49 | CHEMBL242320 |
| BRI-50387 |  | -8.9 | 333.4 | 100 | 0.58 | CHEMBL220806 |
| BRI-50391 |  | -5.8 | 377.4 | 91 | 0.55 | CHEMBL220806 |
| BRI-50392 |  | -5.8 | 318.3 | 97 | 0.51 | CHEMBL242320 |
| BRI-50394 |  | -5.4 | 374.5 | 100 | 0.46 | CHEMBL242320 |
| BRI-50395 |  | -4.2 | 369.3 | 95 | 0.58 | CHEMBL220806 |
| BRI-50453 |  | -30.3 | 458.5 | 91 | 0.52 | CHEMBL345056 |
| BRI-50454 |  | -19.6 | 364.4 | 100 | 0.53 | CHEMBL408261 |
| BRI-50455 |  | -19.7 | 438.5 | 98 | 0.48 | CHEMBL345056 |
| BRI-50456 |  | -21.9 | 408.5 | 100 | 0.49 | CHEMBL345056 |
| BRI-50457 |  | -23.8 | 408.5 | 98 | 0.49 | CHEMBL220806 |
| BRI-50460 |  | -18.4 | 394.4 | 91 | 0.52 | CHEMBL220806 |
| BRI-50461 |  | -22.0 | 446.5 | 100 | 0.49 | CHEMBL242320 |
| BRI-50462 |  | -21.5 | 392.5 | 100 | 0.53 | CHEMBL345056 |
| BRI-50464 |  | -20.0 | 438.5 | 100 | 0.46 | CHEMBL220806 |
| BRI-50465 |  | -30.7 | 472.5 | 99 | 0.53 | CHEMBL345056 |
| BRI-50466 |  | -15.7 | 404.4 | 100 | 0.44 | CHEMBL242320 |
| BRI-50467 |  | -11.4 | 386.5 | 100 | 0.54 | CHEMBL345056 |
| BRI-50469 |  | -19.2 | 406.5 | 100 | 0.52 | CHEMBL345056 |
| BRI-50470 |  | -21.0 | 394.4 | 95 | 0.52 | CHEMBL242320 |
| BRI-50472 |  | -21.7 | 386.3 | 99 | 0.55 | CHEMBL345056 |
| BRI-50473 |  | -22.0 | 408.4 | 100 | 0.51 | CHEMBL242320 |
| BRI-50475 |  | -26.5 | 406.5 | 100 | 0.48 | CHEMBL138867 |
| BRI-50478 |  | -20.7 | 442.5 | 98 | 0.50 | CHEMBL242320 |
| BRI-50479 |  | -26.7 | 460.5 | 97 | 0.52 | CHEMBL242320 |
| BRI-50480 |  | -15.8 | 468.5 | 100 | 0.50 | CHEMBL242320 |
| BRI-50481 |  | -17.5 | 467.6 | 97 | 0.49 | CHEMBL242960 |
| BRI-50482 |  | -26.7 | 386.4 | 100 | 0.45 | CHEMBL242960 |
| BRI-50483 |  | -21.2 | 374.4 | 100 | 0.46 | CHEMBL242960 |
| BRI-50484 |  | -17.5 | 400.4 | 100 | 0.45 | CHEMBL242960 |
| BRI-50486 |  | -21.1 | 436.5 | 95 | 0.47 | CHEMBL242960 |
| BRI-50489 |  | -21.3 | 434.5 | 100 | 0.50 | CHEMBL345056 |
| BRI-50490 |  | -23.3 | 418.5 | 100 | 0.45 | CHEMBL242101 |
| BRI-50493 |  | -32.0 | 400.4 | 95 | 0.43 | CHEMBL242960 |
| BRI-50495 |  | -24.7 | 453.5 | 100 | 0.49 | CHEMBL242960 |
| BRI-50498 |  | -20.0 | 372.4 | 100 | 0.50 | CHEMBL242960 |
| BRI-50499 |  | -20.2 | 437.5 | 90 | 0.54 | CHEMBL345056 |
| BRI-50500 |  | -16.7 | 421.5 | 100 | 0.55 | CHEMBL242320 |
| BRI-50501 |  | -22.0 | 400.4 | 100 | 0.46 | CHEMBL242960 |
| BRI-50502 |  | -23.3 | 505.6 | 91 | 0.58 | CHEMBL345056 |
| BRI-50281 analogs | | | | | | |
| BRI-50401 |  | -30.7 | 410.9 | 100 | 0.48 | CHEMBL3326961 |
| BRI-50404 |  | -29.9 | 449.5 | 90 | 0.57 | CHEMBL428391 |
| BRI-50412 |  | -28.2 | 389.4 | 100 | 0.53 | CHEMBL428391 |
| BRI-50414 |  | -28.1 | 363.4 | 100 | 0.55 | CHEMBL3326961 |
| BRI-50419 |  | -27.6 | 362.4 | 97 | 0.48 | CHEMBL242960 |
| BRI-50425 |  | -26.3 | 349.3 | 91 | 0.55 | CHEMBL3326961 |
| BRI-50426 |  | -26.2 | 365.3 | 95 | 0.57 | CHEMBL3326961 |
| BRI-50428 |  | -25.1 | 403.4 | 91 | 0.57 | CHEMBL3326961 |
| BRI-50431 |  | -24.7 | 349.3 | 99 | 0.57 | CHEMBL3326961 |
| BRI-50432 |  | -24.1 | 377.4 | 96 | 0.55 | CHEMBL3326961 |
| BRI-50434 |  | -23.6 | 364.4 | 100 | 0.56 | CHEMBL259823 |
| BRI-50439 |  | -21.5 | 407.4 | 100 | 0.43 | CHEMBL484413 |
| BRI-50441 |  | -21.3 | 417.4 | 100 | 0.57 | CHEMBL3326961 |
| BRI-50450 |  | -17.1 | 349.3 | 100 | 0.60 | CHEMBL3326961 |
| BRI-50504 |  | -24.1 | 349.3 | 93 | 0.59 | CHEMBL3326961 |
| BRI-50506 |  | -16.3 | 355.4 | 100 | 0.62 | CHEMBL242320 |
| BRI-50507 |  | -15.5 | 355.4 | 99 | 0.59 | CHEMBL242320 |
| BRI-50508 |  | -13.5 | 375.8 | 100 | 0.62 | CHEMBL242320 |
| BRI-50509 |  | -12.4 | 355.4 | 100 | 0.60 | CHEMBL242320 |

**Supplementary Table S5.** Dose-response curves for hit compounds from the cellular PGE_2_ production assay.

| **Cmpd.** | **Structure** | **Dose-response data** | **IC_50_ (nM)** |
| --- | --- | --- | --- |
| ASB-14780 |  | 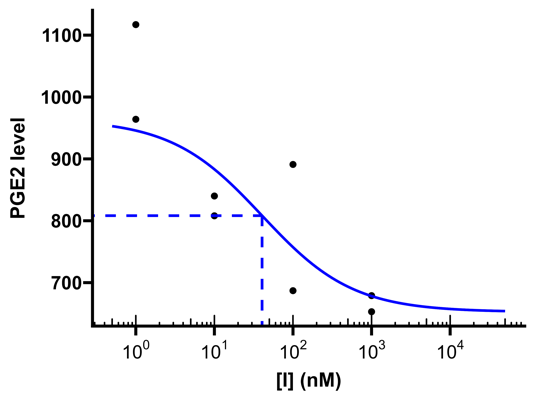 | 40 |
| BRI-50054 |  | 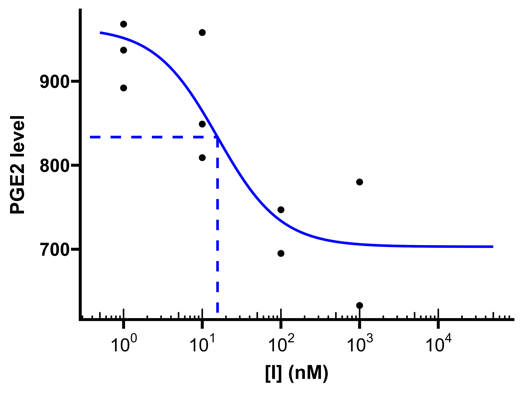 | 16 |
| BRI-50460 |  | 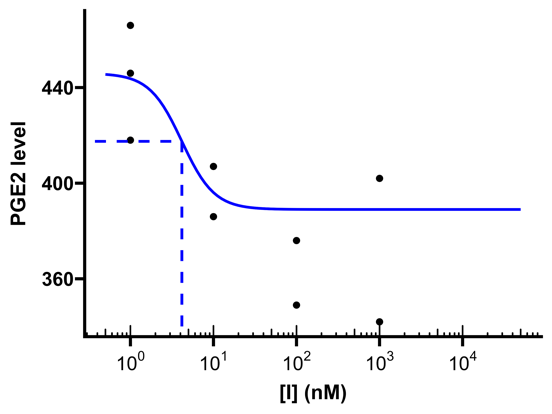 | 4.1 |
| BRI-50469 |  | 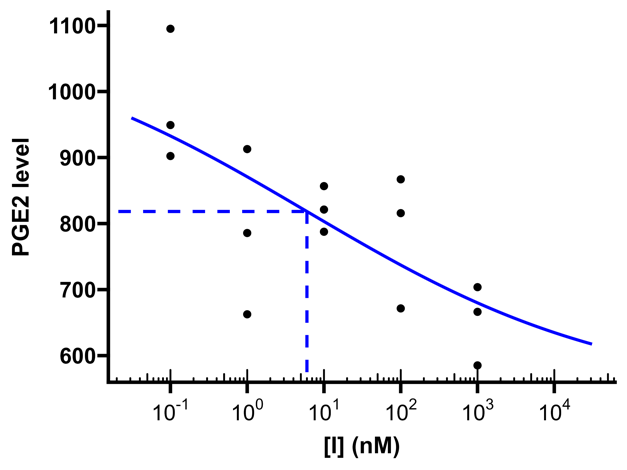 | 6.0 |

**Supplementary Table S6.** LC-MS/MS assay MRM transitions for BRI-50460 pharmacokinetics.

| **Analyte** | **Q1 (*m/z*)** | **Q3 (*m/z*)** | **RT (min)** | **DP (V)** | **CE (V)** | **CXP (V)** | **Internal Standard** |
| --- | --- | --- | --- | --- | --- | --- | --- |
| BRI-50460 | 395.0 | 353.1 | 9.50 | -105 | -31 | -10 | BRI-50469 |
| BRI-50469 | 407.0 | 365.1 | 9.22 | -95 | -33 | -10 | N/A |
